# CRISPR mapping unveils global prevalence of viral predators of Myxobacteria

**DOI:** 10.64898/2026.09.14.751636

**Authors:** Pengwei Li, Jinren Ni

**Affiliations:** Environmental Microbiome and Innovative Genomics Laboratory, College of Environmental Sciences and Engineering, Peking University, Beijing 100871, P. R. China; College of Environmental Sciences and Engineering, Key Laboratory of Water and Sediment Sciences, Ministry of Education, Peking University, Beijing 100871, P. R. China; Eco-environment and Resource Efficiency Research Laboratory, School of Environment and Energy, Peking University Shenzhen Graduate School, Shenzhen 518055, P. R. China

**Keywords:** Myxobacteria, myxophages, viral diversity, host physiology, immune defense

## Abstract

Myxobacteria are social Gram-negative bacteria that perform cooperative predation and form multicellular fruiting bodies, serving as promising biocontrol agents against drug-resistant pathogens. Their associated viruses are therefore important ecological regulators, yet remain largely unexplored. Here, we applied CRISPR spacer-based host-virus linkage analysis to 3,229 myxobacterial genomes and identified 790 nonredundant myxophages spanning diverse global ecosystems. Comparative genomic analyses revealed extensive diversity in the taxonomy, lifestyles, and functional potential of these myxophages. We found that myxophages encode auxiliary metabolic genes involved in carbon, sulfur, iron, and cell-envelope metabolism, which may modulate host physiological processes during infection. Gene-sharing networks, whole-proteome phylogenies, inter-genomic and proteome comparison further identified six previously undescribed myxoviral lineages (MV-1∼MV-6), each represented by complete genomes. Detailed analyses revealed the mechanisms by which these viruses hijack host cellular machinery and reprogram host processes. Notably, MV-1 encodes immune modulators, including the anti-CRISPR protein *AcrIIA15* and a *Cas12m* effector located adjacent to CRISPR arrays, empower myxophages with sophisticated immune evasion and host interference strategies. This study substantially expands the myxoviral diversity and provides insights into virus-host interactions and evolutionary adaptation of myxophages.

**Teaser:** Phages reshape myxobacterial physiology and immune defense.

## Introduction

Predation is a fundamental ecological process that shapes community structure, biodiversity, and nutrient cycling across ecosystems^1,2^. In microbial communities^3,4^, predatory bacteria regulate microbial populations through direct killing and biomass turnover, thereby influencing community assembly and ecosystem function^5,6^. Their ability to eliminate diverse pathogens has also attracted considerable interest for the control of antimicrobial-resistant bacteria^1,7–9^. Among bacterial predators, the best-characterized groups are Bdellovibrio and like organisms (BALOs; phylum Bdellovibrionota)^9,10^ and myxobacteria (phylum Myxococcota)^11,12^. Unlike the obligate predatory BALOs, myxobacteria are facultative predators with a broad prey spectrum encompassing both prokaryotic and eukaryotic microorganisms^9,11^, making them promising biocontrol agents for environmental and biomedical applications^13^.

Myxobacteria are Gram-negative gliding bacteria distinguished by sophisticated multicellular behaviors, including cooperative predation and fruiting body development^12^. They possess some of the largest bacterial genomes, frequently exceeding 10 Mb, which support extensive regulatory networks, secondary metabolism, and complex developmental programs^14^. During predation, myxobacteria employ coordinated “wolf-pack” strategies, secreting extracellular lytic enzymes and bioactive metabolites to kill and digest prey^15^. They also represent one of the richest microbial sources of natural products^16^, including clinically important compounds such as the anticancer drug epothilones^17–21^. Despite more than two centuries of study, their slow growth, complex life cycle, and demanding cultivation requirements have limited the availability of cultured isolates^20,22,23^, leaving much of their diversity and ecology unexplored^11,24^.

Viruses are major drivers of microbial evolution, mortality, metabolism, and biogeochemical cycling^25,26^. Yet, compared with other bacterial lineages, the virome associated with myxobacteria remains largely unexplored, primarily because of the scarcity of cultured hosts. Current knowledge is largely confined to phages infecting model organisms such as *Myxococcus xanthus*^27–29^, leaving the diversity, phylogenetic breadth, functional capacity, and infection mechanisms of myxophages largely unknown. This knowledge gap limits our understanding of how viruses influence predatory bacteria and their potential applications as living biocontrol agents^1,8^.

Advances in metagenomics and clustered regularly inter-spaced short palindromic repeat (CRISPR)^30^ spacer-based virus discovery have enabled the identification of viruses infecting uncultivated microorganisms^31,32^, including Candidate Phyla Radiation (CPR) bacteria^33^, Asgard archaea^34,35^, DPANN archaea^36^, anaerobic methanotrophs (ANME)^37^, and methanogenic archaea^38^. Building on these advances, we systematically characterized CRISPR–Cas systems in globally distributed myxobacterial genomes and used CRISPR-guided host–virus linkage to reconstruct the myxobacterial virome. This study expands the known diversity of myxophages and provides insights into their evolutionary relationships, functional repertoires, and interactions with their bacterial hosts, establishing a foundation for understanding the ecological roles of myxophages and their potential contributions to myxobacteria-based biocontrol.

## Results

### Overview of biosynthetic gene clusters, antimicrobial peptides, defense systems, secretion systems and the mobilome of myxobacteria

We curated 3,229 myxobacterial genomes from the National Center for Biotechnology Information (NCBI) GenBank and six publicly available datasets spanning diverse ecosystems, including marine, soil, freshwater, engineered, rhizosphere, host-associated, glacier, and extreme ecosystems (Fig. S1, Supplementary Table 1, see Methods). Genome size and GC content varied substantially among classes within the phylum Myxococcota (Fig. S1B). However, the available genomes were taxonomically uneven, with most belonging to the orders Polyangiales and Myxococcales (Fig. S1C), highlighting the limited genomic representation of many myxobacterial lineages and future sequencing efforts are needed.

To characterize the functional repertoire underlying the distinctive biology of myxobacteria^39^, we systematically identified biosynthetic gene clusters (BGCs), antimicrobial peptides (AMPs), anti-phage defense systems (DSs), and secretion systems (TXSSs) across all genomes (Fig. 1, Supplementary Table 2-5). In total, we detected 44,698 BGCs (13.8 per genome), 2,334 AMPs (0.7 per genome), 21,007 defense systems (6.5 per genome), and 5,130 secretion systems (1.6 per genome) using strict filtering standards (see Methods). Myxobacterial genomes encoded a higher abundance of defense systems than the average reported for RefSeq prokaryotic genomes^40^. Restriction–modification (R–M), CRISPR–Cas, Septu, and AbiE were the dominant defense systems (Fig. 1A, H), consistent with previous studies^41^. Type II secretion systems (T2SSs) were the most prevalent secretion systems (Fig. 1B, E), consistent with their established role in extracellular slime secretion^39^. Type IVa pili (T4aP) are cell surface filaments important for surface motility, adhesion to surfaces, DNA uptake, biofilm formation, and virulence^42,43^. These functions are especially important for myxobacteria as its special lifestyle^42^. T4aP were absent or markedly reduced in several classes, including UBA9042, UBA727, UBA796, and Polyangia (Fig. 1E), indicating substantial differences in surface-associated lifestyles among myxobacterial lineages. The widespread occurrence of BGCs and AMPs further underscores the exceptional biosynthetic potential of myxobacteria^16^ (Fig. 1C, D, F, G). Bacteria rapidly adapt to fluctuating environments through horizontal gene transfer (HGT)^44^, facilitated by mobile genetic elements (MGEs)^45^, which are broadly classified into plasmids, bacteriophages (phages), and integrative elements^46^, including insertion sequences (ISs), transposons (Tn)^47^, integrons, and integrative conjugative/mobilizable elements (ICEs/IMEs)^48^. To explore the contribution of MGEs to myxobacterial functional traits, we systematically identified plasmids, prophages, insertion sequences (ISs), transposons (Tns), integrons, and integrative conjugative/mobilizable elements (ICEs/IMEs). A total of 11,969 MGEs were detected (Fig. 1I, Supplementary Table 6-10), comprising 5,770 plasmids, 545 prophages, 5,336 ISs, 192 integrons, 116 composite transposons (Tns), and 10 ICEs/IMEs. These MGEs carried diverse functional cargo, including defense systems, secretion systems, and AMPs (Fig. 1J–L). Plasmids were the principal vectors for all three functional categories (Fig. 1J, Fig. S2), whereas prophages preferentially encoded defense systems, particularly R– M, AbiE, and SoFIC. Notably, plasmids were enriched in pT4SSt rather than the more abundant chromosomal T2SS, suggesting functional specialization in the dissemination of secretion-associated traits^49^.

**Fig. 1.**
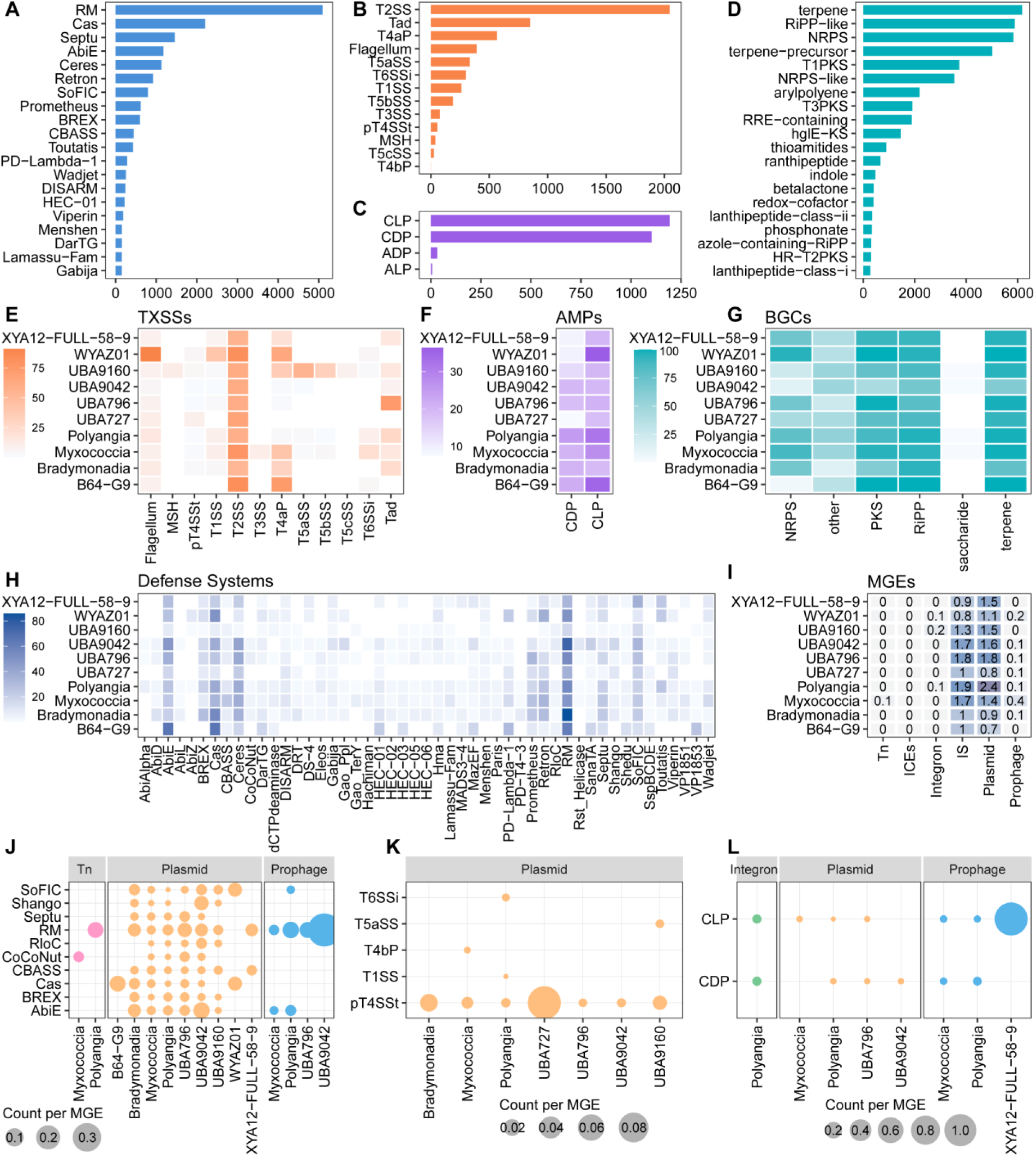
Functional repertoire and mobilome-associated genetic traits of myxobacteria. **(A)** Distribution of the 20 most abundant anti-phage defense systems (DSs) identified across myxobacterial genomes. **(B)** Distribution of secretion systems (TXSSs) identified in myxobacterial genomes. **(C)** Distribution of antimicrobial peptides (AMPs). CLP, cationic linear peptide; CDP, cationic disulfide peptide; ADP, anionic disulfide peptide; ALP, anionic linear peptide. **(D)** Distribution of biosynthetic gene clusters (BGCs). **(E–H)** Prevalence of TXSSs, AMPs, BGCs, and DSs, respectively, across the ten most represented myxobacterial classes. Heat-map colour intensity indicates the proportion of genomes within each class encoding the corresponding feature. **(I)** Distribution of mobile genetic elements (MGEs) across the ten most represented myxobacterial classes. MGE abundance is normalized to the number of genomes in each class. **(J–L)** Functional genes carried by MGEs, including DSs (J), TXSSs (K), and AMPs (L). Dot size is proportional to the number of genes normalized to the corresponding number of MGEs.

To establish a CRISPR spacer database for subsequent virus identification, we systematically characterized CRISPR–Cas systems in the 3,229 myxobacterial genomes. After quality filtering and manual curation (see Methods), 474 genomes were found to contain complete or nearly complete CRISPR–Cas loci (Fig. 2). These systems comprised two classes, four types, and thirteen subtypes (Supplementary Table 11). Class 1 systems predominated, with type I (I-A, I-B, I-C, I-E, I-F, and I-G) and type III (III-A, III-B, III-C, and III-D) identified in 470 genomes, whereas class 2 systems, including subtypes II-A, V-M, and V-B1, were detected in only eight genomes (Fig. 2B). Consistent with previous studies^50^, type I and type III systems dominated the CRISPR–Cas repertoire of myxobacteria (Fig. 2A). The results provide a comprehensive overview and enrich our knowledge of the diversity of CRISPR–Cas system classes, types and subtypes across myxobacterial taxa. It should be noted that several genomes contained CRISPR arrays but lacked complete Cas operons, precluding confident subtype assignment.

**Fig. 2.**
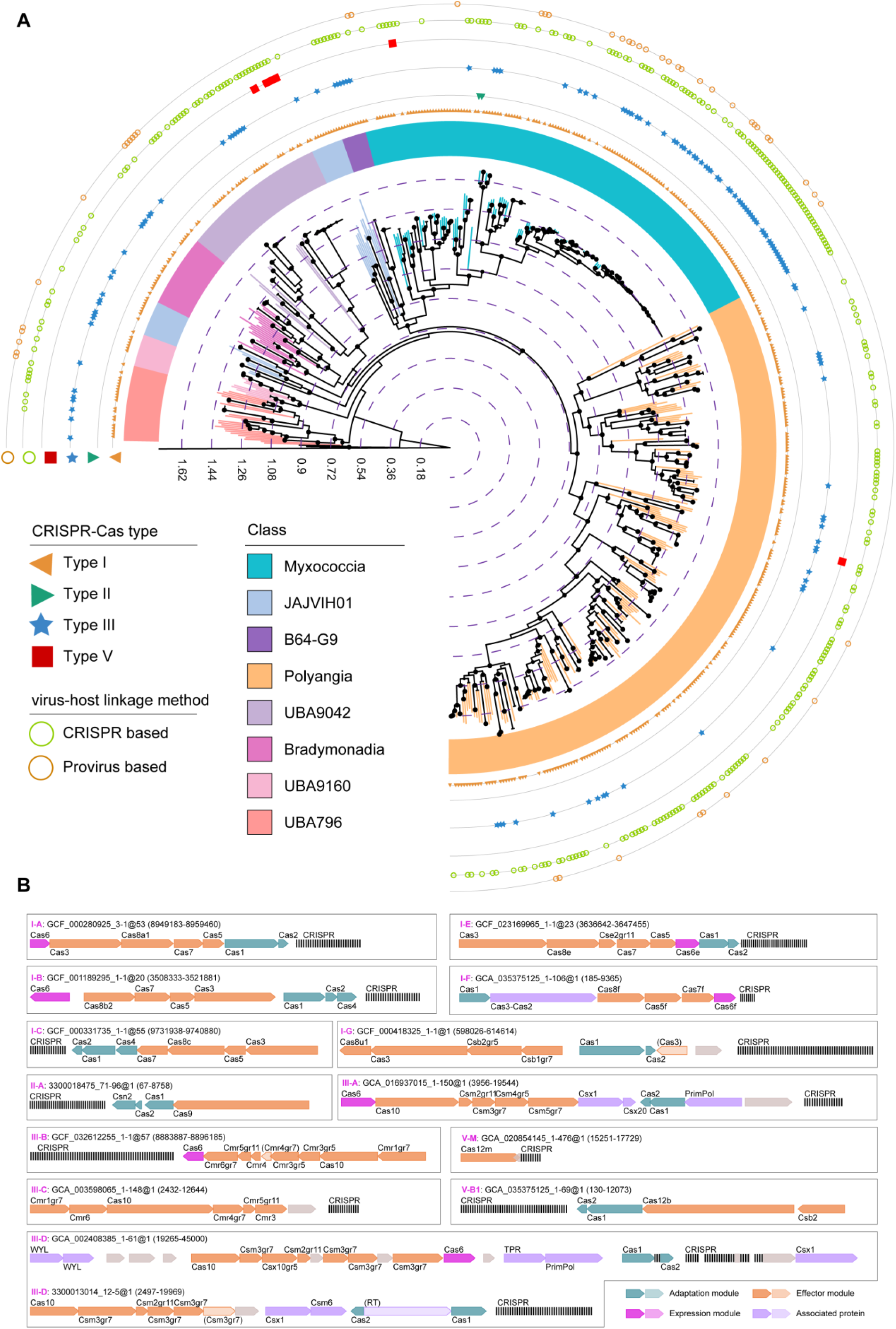
Diversity and phylogenetic distribution of CRISPR–Cas systems in myxobacteria. **(A)** Maximum-likelihood phylogeny of 565 genomes containing identifiable CRISPR–Cas systems or predicted to serve as hosts of the recovered viruses. The tree is infered from 120 conserved bacterial marker genes and rooted with three Cyanobacteriota genomes. The two outermost rings indicate the methods used for virus-host assignment. Inner rings denote the CRISPR–Cas types and subtypes identified in each genome. Black circles indicate branches with bootstrap support >90%. **(B)** Representative genetic organization of each identified CRISPR–Cas subtype. One representative locus is shown for subtypes detected in multiple genomes. Gene arrows indicate the organization and orientation of CRISPR-associated (cas) genes and adjacent CRISPR arrays. CRISPR–Cas subtypes are labeled within each panel.

### Expansive diversity of viruses associated with myxobacteria

To identify viruses associated with myxobacteria, we integrated viral sequences from the IMG/VR v4 database^51^, the VIRE database^52^, Virus-Host DB^53^ (accessed on 8 December, 2025), and the collection of myxobacterial genomes using CRISPR spacer-protospacer matching and prophage detection (see Methods). This approach recovered 1,944 viral sequences through CRISPR spacer targeting and 51 prophages from myxobacterial genomes. In addition, five previously reported reference myxophages (Mx8, Mx4, Mx9, Mx1, Mx4 ts27htf-1hrm-1)^27,50,54^ were retrieved from Virus-Host DB. In total, 2,000 viral sequences were clustered into 1,112 nonredundant viral operational taxonomic units (vOTUs). Following stringent quality filtering (see Methods), 790 high-confidence vOTUs were retained to establish the Myxobacterial Virus Database (MVD, Fig. 3A, Supplementary Table 12).

**Fig. 3.**
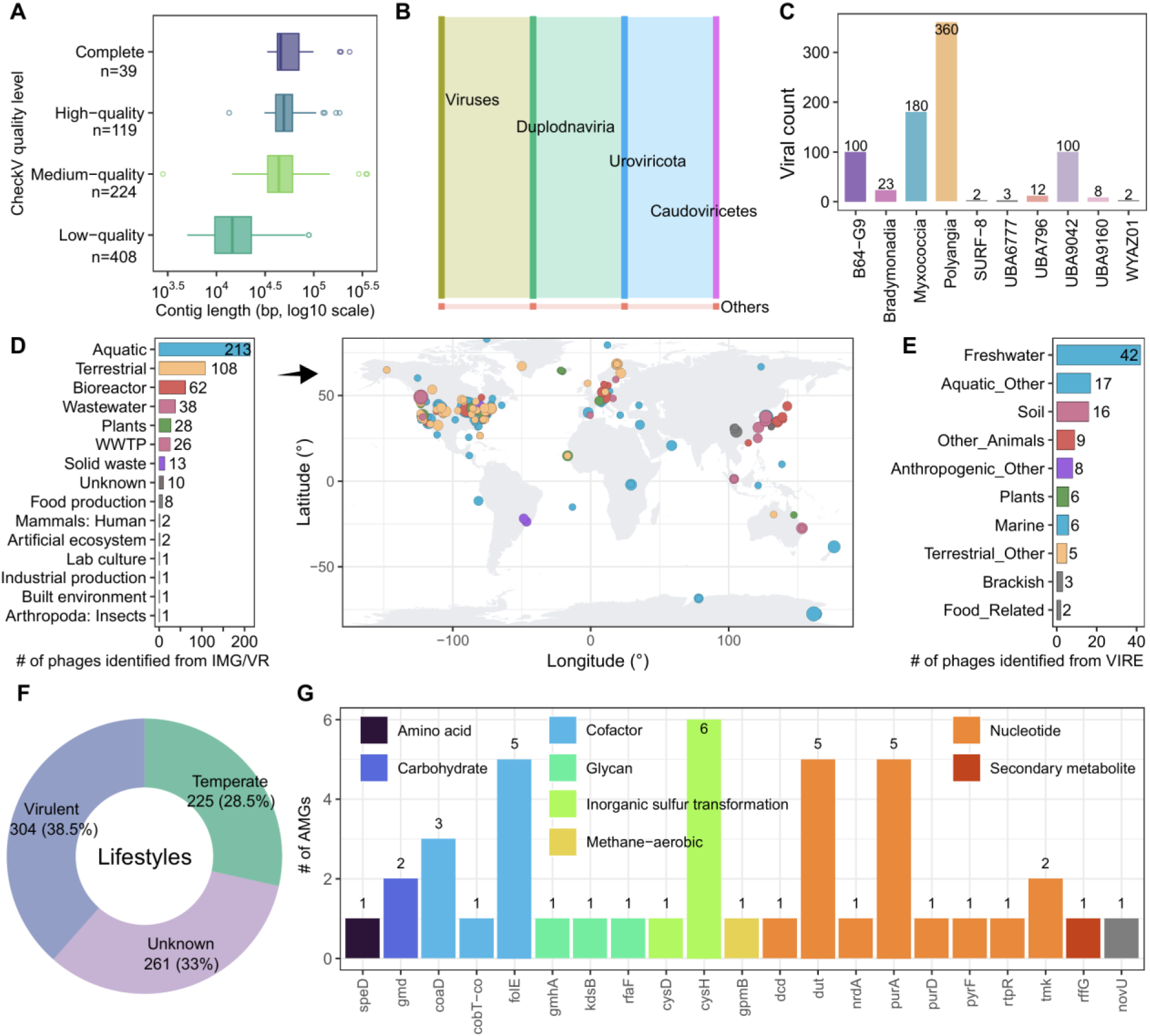
Global diversity, ecological distribution, and metabolic potential of myxobacterial viruses. **(A)** Quality assessment of the recovered myxobacterial viral genomes based on CheckV. **(B)** Taxonomic classification of the retained viral operational taxonomic units (vOTUs). **(C)** Taxonomic distribution of the predicted hosts at the class level. **(D)** Ecosystem origins and geographic distribution of vOTUs recovered from the IMG/VR database. Circles on the map indicate sampling locations, with circle size proportional to the number of vOTUs recovered from each site. Colours denote ecosystem types, and the accompanying bar chart summarizes the number of vOTUs recovered from each ecosystem. vOTUs lacking ecosystem or geographic metadata are not shown here. **(E)** Ecosystem distribution of vOTUs recovered from the VIRE database. **(F)** Predicted viral lifestyles, classified as virulent, temperate, or unknown. (**G**) High-confidence auxiliary metabolic genes (AMGs) involved in carbon, nitrogen, sulfur, and phosphorus metabolism. Gene names follow the Kyoto Encyclopedia of Genes and Genomes (KEGG) nomenclature.

Taxonomic classification showed that 777 (98.4%) vOTUs belonged to *Caudoviricetes* (Fig. 3B). Comparative genomic analyses further revealed distinct genomic and proteomic characteristics among phages infecting different myxobacterial lineages, including GC content, average protein molecular weight, carbon-, nitrogen-, and sulfur-atoms per residue side chain (C/N/S-ARSC), and amino acid usage patterns (Fig. 3C, Fig. S3), suggesting adaptation to host-specific genomic and metabolic environments. Few viral genomes were recovered for the classes Bradymonadia, SURF−8, UBA6777, UBA796, UBA9160, and WYAZ01 (Fig. 3C), likely reflecting the limited availability of host genomes (Fig. S1) or the low prevalence of CRISPR–Cas systems in these lineages, which constrained spacer-guided viral discovery. These limitations suggest that substantial viral diversity associated with these taxa remains undiscovered and warrants further investigation. Most viral genomes originated from aquatic and terrestrial environments (Fig. 3D, E), consistent with the ecological distribution of myxobacteria (Fig. S1A). Lifestyle prediction classified 38.5% of myxophages as virulent and 28.5% as temperate, whereas 33% could not be confidently assigned (Fig. 3F).

To investigate the metabolic potential of the myxobacterial virome, we profiled auxiliary metabolic genes (AMGs) involved in carbon, nitrogen, sulfur, and phosphorus metabolism (Fig. 3G). The recovered myxophages encoded a diverse repertoire of AMGs spanning multiple metabolic pathways, suggesting their potential to reprogram host metabolism during infection. Genes involved in nucleotide metabolism (e.g., *dut* and *purA*) and cofactor biosynthesis (predominantly *folE*) were abundant, indicating that myxophages may enhance nucleotide biosynthesis and metabolic capacity to support efficient viral replication. Sulfur metabolism was represented primarily by the phosphoadenylyl-sulfate reductase gene (*cysH*), highlighting sulfur assimilation as a prominent auxiliary function. In addition, we identified AMGs involved in carbohydrate and glycan metabolism, including *gmd*, *gmhA*, *kdsB*, and *rfaF*, which are associated with the biosynthesis and modification of cell-surface polysaccharides^55^. These genes suggest that myxophages may alter the composition of the myxobacterial cell envelope during infection, potentially influencing phage susceptibility, superinfection dynamics^56^, and cell-cell interactions that underpin the social lifestyle of myxobacteria^57^.

### Identification of novel viral groups of myxobacteria

To assess the novelty of the recovered viruses, all nonredundant viral operational taxonomic units (vOTUs) were compared with previously reported myxophages^27,28,58^ and reference prokaryotic viruses from the NCBI RefSeq database using gene-sharing network analysis (Fig. 4A). Notably, six viral clusters, each represented by at least one complete genome, showed no detectable connections to any reference viruses, suggesting that they represent previously uncharacterized lineages within the myxobacterial virosphere. Proteome-based phylogenetic analysis further placed the representative genomes of these clusters on six well-separated branches distinct from known viruses (Fig. 4B). Consistent with their phylogenetic divergence, these viruses shared <70% intergenomic similarity, below the VIRIDIC genus-level threshold^59^ and <10% orthologous proteins with reference viruses or between clusters (Fig. 4C, D). Comparative genome analysis further revealed conserved gene synteny within each cluster but substantial variation among clusters (Fig. 4E). Typically, viral family-level assignments in proteome-based phylogenies are made when branch lengths exceed 0.05^36,38^, and viruses from different families usually share less than 10% of orthologous proteins^36,38^. These independent lines of evidence support the proposal of six novel myxobacterial viral groups (MVs): MV-1 (GCA_020854145_1-476), MV-2 (IMGVR_UViG_3300014208_000007), MV-3 (IMGVR_UViG_3300043313_001413), MV-4 (vire_genome_817650), MV-5 (vire_genome_735416), and MV-6 (vire_genome_1676178) (Supplementary Table 13; see Methods), which likely represent founding members of previously undescribed viral lineages.

**Fig. 4.**
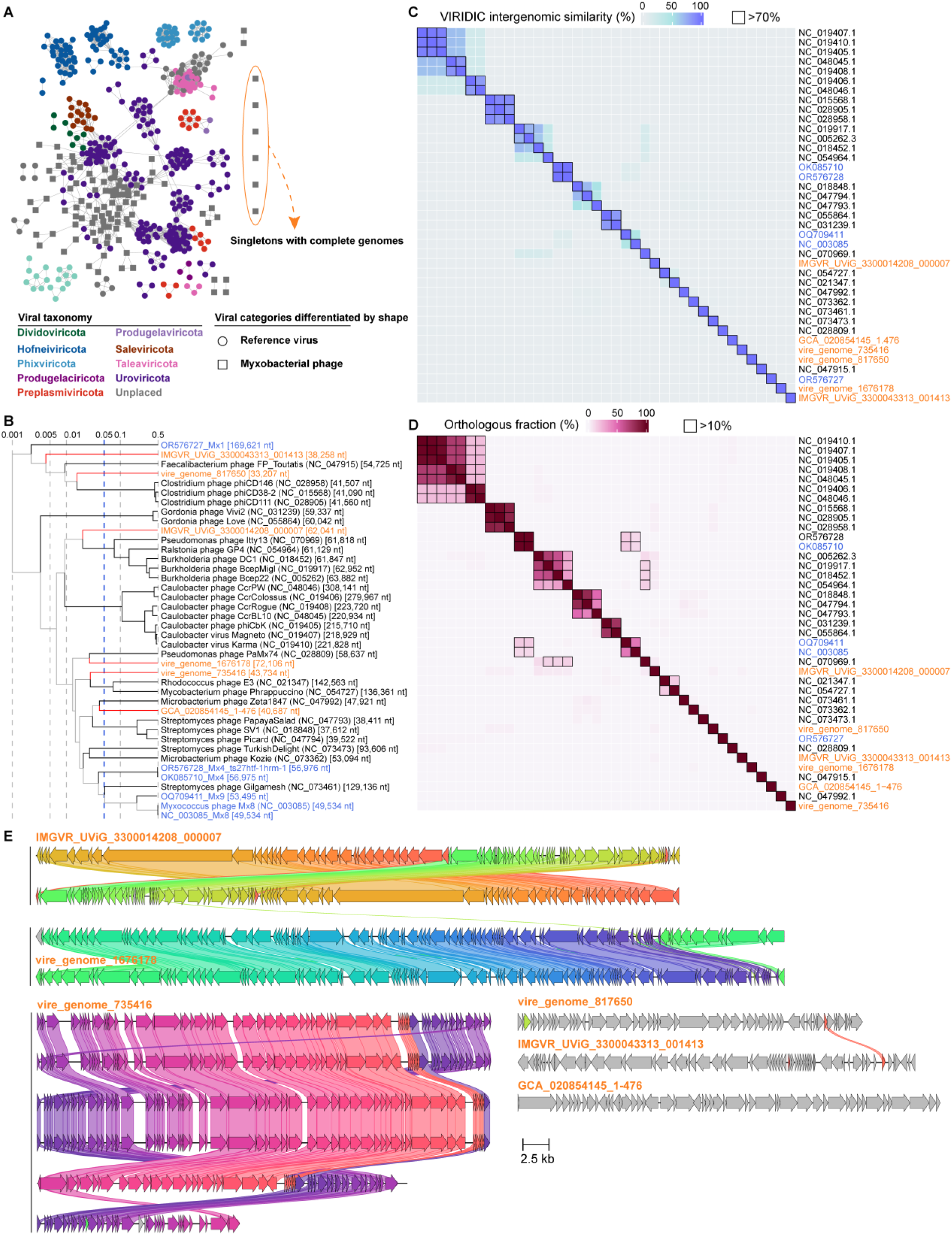
Genome-based taxonomic assignment of proposed novel viruses. **(A)** Gene-sharing network of viruses and reference viruses. Each node represents a viral genome, and edges indicate shared protein clusters. Node colors denote viral taxonomy, and node shapes distinguish virus categories. Reference viruses were retrieved from the NCBI Viral RefSeq database and previously reported myxophages (see Methods). Orange circles highlight the six newly proposed viral groups according to the gene-sharing network (Supplementary Table 13). **(B)**Whole-proteome phylogeny of the proposed viral groups and representative reference viruses. Newly proposed viral groups are shown in orange, previously reported myxobacterial phages in blue, and reference viruses in black. The dashed blue line indicates the proposed family-level branch-length threshold (0.05). **(C)** Pairwise intergenomic nucleotide similarity among the proposed viral groups and reference viruses, calculated using VIRIDIC. Values >70%, corresponding to the ICTV genus-level threshold, are outlined in black. **(D)** Pairwise orthologous protein fractions shared among viral genomes. Values >10% are outlined in black. **(E)** Genomic comparison of the six proposed viral groups. Homologous proteins are connected and coloured consistently across genomes. Shading intensity indicates amino acid sequence identity (>30%). Complete viral genomes are highlighted in orange fonts for their IDs.

All six proposed viral groups belong to the class *Caudoviricetes* within the realm Duplodnaviria (Supplementary Table 13) and encode the conserved hallmark proteins of head-tailed bacteriophages, including HK97-like major capsid proteins, terminase large subunits, portal proteins, and tail structural proteins (Fig. 5, Fig. S4-5, and Supplementary Table 14). MV-2 (from groundwater), MV-3 (from soil), and MV-4 (from gut) each contained a protospacer targeted by CRISPR spacers recovered from the same six Myxococcota genomes identified in activated sludge from a wastewater treatment plant, suggesting a shared host association despite their distinct environmental origins. The genome size and GC content of MV-3 (38.2 kb, 50.7%) and MV-4 (33.2 kb, 45.9%) is smaller than that of MV-2 (62.0 kb, 60.11%). MV-5 and MV-6, both recovered from soil, were each targeted by spacers derived from members of the order Polyangiales, whereas the host of MV-1, recovered from biofilm in wastewater treatment plant, was assigned to Polyangiales based on blastp against the NCBI NR database (see Methods). Most of these viruses and their predicted hosts originated from soil or engineered environments, with only one recovered from an aquatic habitat, consistent with the habitat preference of myxobacteria^60^. The high microbial density characteristic of bioreactors may increase opportunities for interactions among myxobacteria, their prey, and associated viruses, potentially facilitating virus-host encounters and viral transmission^61,62^.

**Fig. 5.**
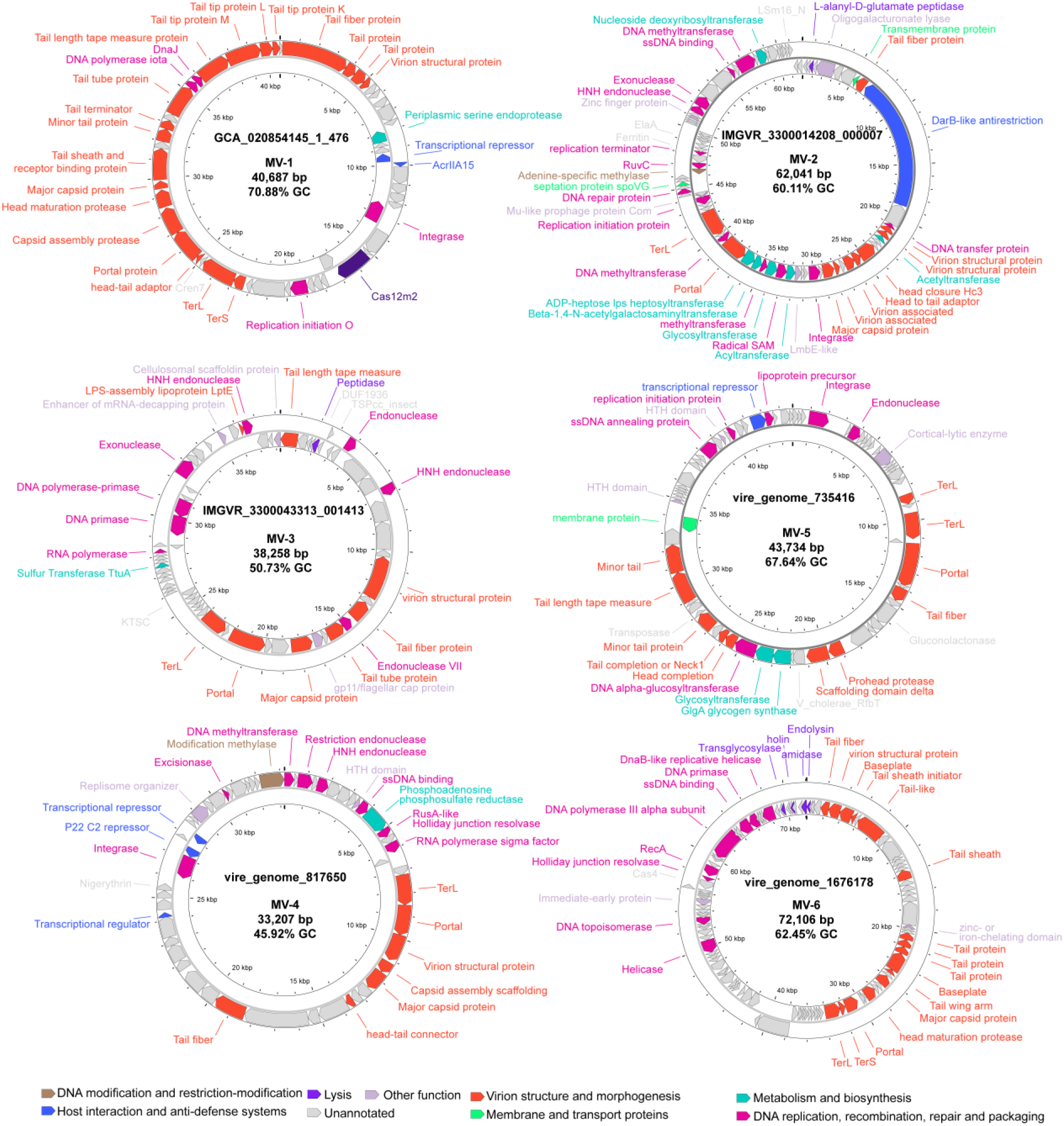
Genomic architecture and functional annotation of the proposed myxobacterial viruses. Complete genomes are identified by the presence of direct terminal repeats (DTRs) or by circular genome topology, as indicated. Gene arrows represent predicted open reading frames and their transcriptional orientation. Colours denote predicted functional categories. TerL, terminase large subunit; TerS, terminase small subunit. Detailed functional annotations are provided in Supplementary Table 14.

### Replication, transcription, and translation features of the proposed MVs

The six newly identified MVs encode diverse functional potentials associated with genome replication, transcriptional regulation, and protein synthesis. Many of these proteins, including DNA helicases and DNA primases, are widely conserved across viruses and cellular organisms, reflecting the exploitation of fundamental host molecular processes during infection. Nevertheless, a large proportion of predicted open reading frames could not be assigned functions using current public databases (Fig. 5, see Methods and Supplementary Table 14), highlighting the substantial unexplored functional diversity of myxophages.

BACPHILIP^63^ predicted MV-1, MV-2, MV-4, and MV-5 to be temperate phages, whereas MV-3 and MV-6 were classified as virulent. Consistent with these predictions, all four temperate viral groups encoded integrases^64^, together with additional lysogeny-associated proteins, including replication initiation proteins, transcriptional repressors^64,65^, and excisionases^64^, which collectively regulate the transition between lysogenic and lytic lifestyles. MV-4 additionally encoded a RuvC Holliday junction resolvase, which may facilitate the resolution of recombination intermediates during prophage integration and excision^66^. In contrast, no canonical temperate-associated marker genes were identified in MV-3 or MV-6.

Genes involved in DNA replication and recombination were widespread across the proposed viral groups. Exonucleases (MV-2 and MV-3), endonucleases (MV-2, MV-3, MV-4, and MV-5), and single-stranded DNA-binding proteins (MV-2, MV-4, MV-5, and MV-6) were frequently identified, suggesting conserved mechanisms supporting viral DNA replication and genome maintenance^67^. HNH endonucleases, encoded by MV-2, MV-3, and MV-4, may participate in DNA cleavage during genome maturation and packaging in concert with the terminase large subunit and portal protein^68^. Several viruses also encoded DNA polymerases^69^, primases, DNA helicases^70^, or polymerase–primase fusion proteins^71^, indicating an expanded capacity to support viral genome replication. Consistent with their predicted lytic lifestyles, both virulent viruses encoded proteins involved in host cell lysis. MV-3 encoded a peptidase, whereas MV-6 possessed a more complete lysis module comprising holin, endolysin, amidase, and transglycosylase, suggesting coordinated degradation of the bacterial cell envelope to enable progeny virion release^72^.

Beyond genome replication, the proposed myxophages encode proteins that may modulate host transcriptional and translational processes^73^ (Fig. 5). MV-3 carries an enhancer of mRNA-decapping protein, which has been reported to promote host mRNA degradation, thereby suppressing host gene expression and potentially redirecting translational resources toward viral protein synthesis^74^. Additional transcription-associated proteins, including RNA polymerase subunits (MV-3 and MV-4) and a transcriptional regulator (MV-4), were also identified. Translation-associated proteins were less common but included the molecular chaperone DnaJ in MV-1, which facilitates protein folding^75,76^, and the tRNA sulfurtransferase TtuA in MV-3, which catalyzes sulfur modification of tRNA^77^ and may contribute to efficient and accurate translation during infection.

### Viral genes potentially enhance host metabolism and physiological functions

The proposed myxophages encode diverse auxiliary metabolic genes (AMGs), indicating the potential to modulate host metabolism beyond processes directly required for viral genome replication and particle assembly^78^. Several AMGs are associated with carbon metabolism, sulfur assimilation, iron homeostasis, and cell-envelope biogenesis (Fig. 5). For example, the glycogen synthase gene (*glgA*) encoded by MV-5 may redirect carbon flux toward glycogen biosynthesis^79^, potentially altering intracellular carbon allocation during infection. Likewise, phosphoadenosine phosphosulfate reductase (*cysH*, encoded by MV-4)^80^ and the tRNA sulfur transferase (*ttuA*, encoded by MV-3)^81^ are involved in sulfur metabolism, which supports cysteine biosynthesis, Fe–S cluster assembly, and cellular redox balance. MV-2 additionally encodes ferritin^82^, whereas MV-4 encodes the putative antioxidant protein nigerythrin^83^, suggesting potential roles in iron homeostasis and oxidative stress mitigation during infection. Such metabolic functions may be particularly advantageous in myxobacteria, whose predatory lifestyle and extensive secondary metabolism impose substantial energetic and physiological demands^84^. We also identified multiple genes involved in lipopolysaccharide and glycan biosynthesis, suggesting that myxophages may modify the composition of the myxobacterial cell surface. Such remodeling could influence phage receptor accessibility^85^, alter susceptibility to primary or secondary phage infection^86^, and affect cell–cell interactions underlying gliding motility, cooperative predation, and multicellular development^87^.

Collectively, these AMGs extend beyond housekeeping metabolic functions and appear well aligned with the distinctive physiology of predatory myxobacteria. Cooperative behaviors such as gliding motility, multicellular aggregation, extracellular enzyme secretion, and prey degradation require substantial metabolic investment^12,23,54,84^. Viral modulation of sulfur metabolism, iron homeostasis, carbon allocation, and cell-envelope composition may therefore help maintain host physiological activity while promoting efficient viral replication. Although these functional assignments are supported by gene annotation and comparative genomics, experimental validation using tractable virus–host systems, together with transcriptomic and metabolomic analyses, will be required in future studies.

### Distinctive interactions between the proposed MVs and their hosts

In addition to metabolic genes, the proposed myxophages encode several proteins that may facilitate interactions with host defense systems (Fig. 5, Fig. S6). DNA methyltransferases encoded by MV-2 and MV-4 suggest the potential to methylate viral DNA, thereby reducing susceptibility to host restriction–modification (R–M) systems^88,89^. Interestingly, MV-4 also encodes a restriction endonuclease together with a cognate methyltransferase, forming a complete R–M system that may function as a phage-encoded defense module to limit invasion by competing mobile genetic elements^86^. MV-2 and MV-5 each encode glycosyltransferases that may catalyze glycosylation of DNA, proteins, or lipids^90^. Viral glycosylation has been implicated in protection against host antiviral defenses and in limiting superinfection^90^, suggesting similar functions in myxophages. We also identified the anti-CRISPR protein AcrIIA15 in MV-1, indicating a potential mechanism for evading CRISPR–Cas-mediated immunity^40,91^ (Fig. S7).

CRISPR–Cas systems are adaptive immune mechanisms widespread in prokaryotes but rarely found in viruses^92^. A particularly notable finding was the identification of an atypical type V-M CRISPR–Cas system in the complete genome of MV-1 (Fig. 6A). This locus comprises a Cas12m effector (designated MvCas12m) and an adjacent CRISPR array^93,94^ but lacks the adaptation genes (*cas1*, *cas2*, and *cas4*)^94^, consistent with previously described virus-encoded CRISPR–Cas systems^36,95^. Unlike canonical type V systems, no recognizable tracrRNA was identified. Furthermore, the CRISPR repeats share only 58% sequence identity with those of the predicted host, making acquisition of spacers directly from the host CRISPR machinery appear unlikely (Fig. 6C). Alternative evolutionary scenarios, including *de novo* expansion of repeat sequences (ωRNA segments) or transposon-associated acquisition^96^, therefore merit consideration but require experimental validation. The array contains one consensus direct repeat and eight spacers, although no confident protospacer targets were identified (see Methods).

**Fig. 6.**
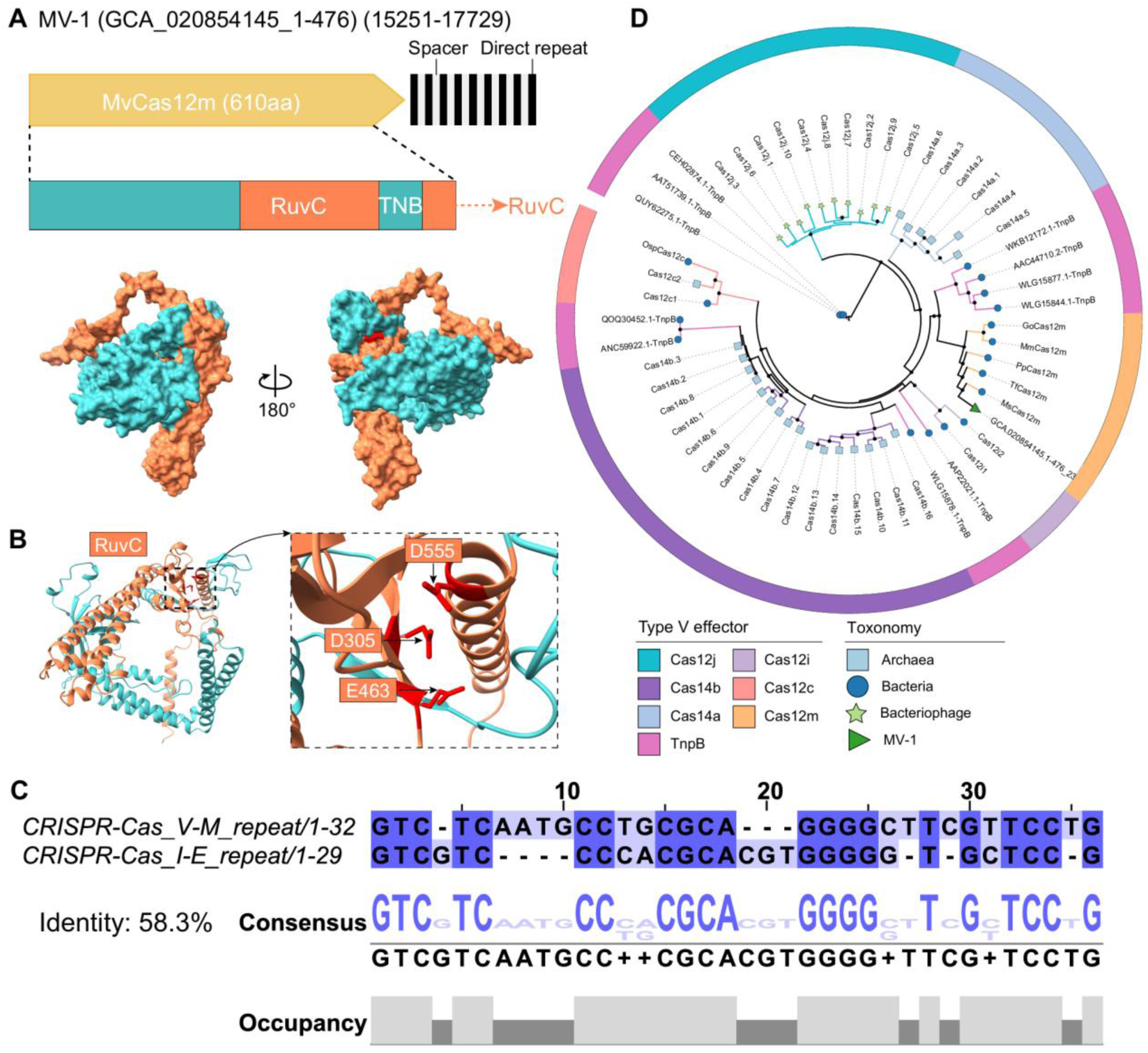
The virus-encoded type V CRISPR–Cas system identified in MV-1. **(A)** Genomic organization of the type V-M CRISPR–Cas locus encoded by MV-1 and the AlphaFold3- predicted structure of the Cas12m effector (MvCas12m). Numbers in brackets indicate the genomic coordinates of the CRISPR–Cas locus within the MV-1 genome. **(B)** AlphaFold3-predicted structure of MvCas12m. The inset highlights the catalytic residues of the RuvC nuclease domain (red). **(C)** Local pairwise alignment of CRISPR repeat sequences from the viral type V-M system and the corresponding host type I-E CRISPR–Cas system. **(D)** Maximum-likelihood phylogeny of type V CRISPR effectors and TnpB nucleases. Black dots indicate branches with bootstrap support >70%. Cas14a (i.e., Cas12f1); Cas14b (i.e., Cas12f2).

Virus-encoded mini-CRISPRs have been previously identified in bacteriophages and archaeal viruses, and they may serve to exclude superinfection by other viruses^36,95,97,98^. MvCas12m is a compact effector protein (610 amino acids, 69.01 kDa) that remains the conserved RuvC catalytic residues (D305, E463, and D555) (Fig. 6B and Fig. S8), suggesting that it may possess nuclease activity comparable to other Cas12m proteins^94^. To our knowledge, this represents the first Cas12m identified in a bacteriophage, whereas previously characterized Cas12m homologs have been restricted to prokaryotic genomes. Phylogenetic analysis further suggests that MvCas12m was likely acquired through horizontal gene transfer from an uncultured bacterial lineage (Fig. 6D). The presence of separate TnpB gene within different CRISPR effector clades implies multiple independent evolutionary trajectories from TnpB to type V CRISPR effectors, as discussed by a recent study^99^. Because virus-encoded CRISPR– Cas systems can suppress competing mobile genetic elements or modulate host functions, the MV-1 V-M CRISPR–Cas system may contribute to viral persistence by mediating interactions with the host and competing viruses^95,100,101^.

## Discussion

This study provides a global view of the previously unexplored virome associated with myxobacteria and reveals diverse virus-host interactions in this ecologically important group of predatory bacteria. By combining CRISPR-guided host assignment with metagenomic analyses, we identified 790 high-confidence myxophages, including six previously undescribed viral groups, substantially expanding the known diversity of myxobacterial viruses. Their broad distribution across freshwater, terrestrial, and engineered environments mirrors the ecological distribution of myxobacteria^60^ (Fig. S1), whereas the limited number of complete viral genomes suggests that the currently recovered diversity represents only a fraction of the natural myxobacterial virome.

The recovered myxophages encode diverse functional modules involved in viral replication, host interaction, and metabolic regulation^64,72,90^ (Fig. 7). In addition to conserved replication and structural proteins^64,102^, we identified genes associated with immune evasion, including DNA methyltransferases^103^ and the anti-CRISPR protein AcrIIA15^104,105^, as well as auxiliary metabolic genes related to carbon allocation, sulfur metabolism, iron homeostasis, and cell-envelope modification^68,76,88,90,104,106,107^. These findings suggest that myxophages may influence host physiology beyond viral propagation alone^108^. We also identified a virus-encoded type V-M CRISPR–Cas system containing a Cas12m effector, representing, to our knowledge, the first report of a Cas12m protein encoded by a bacteriophage^96^. Together, these observations expand the repertoire of virus-encoded immune and metabolic functions associated with bacterial viruses^93,96,107^.

**Fig. 7.**
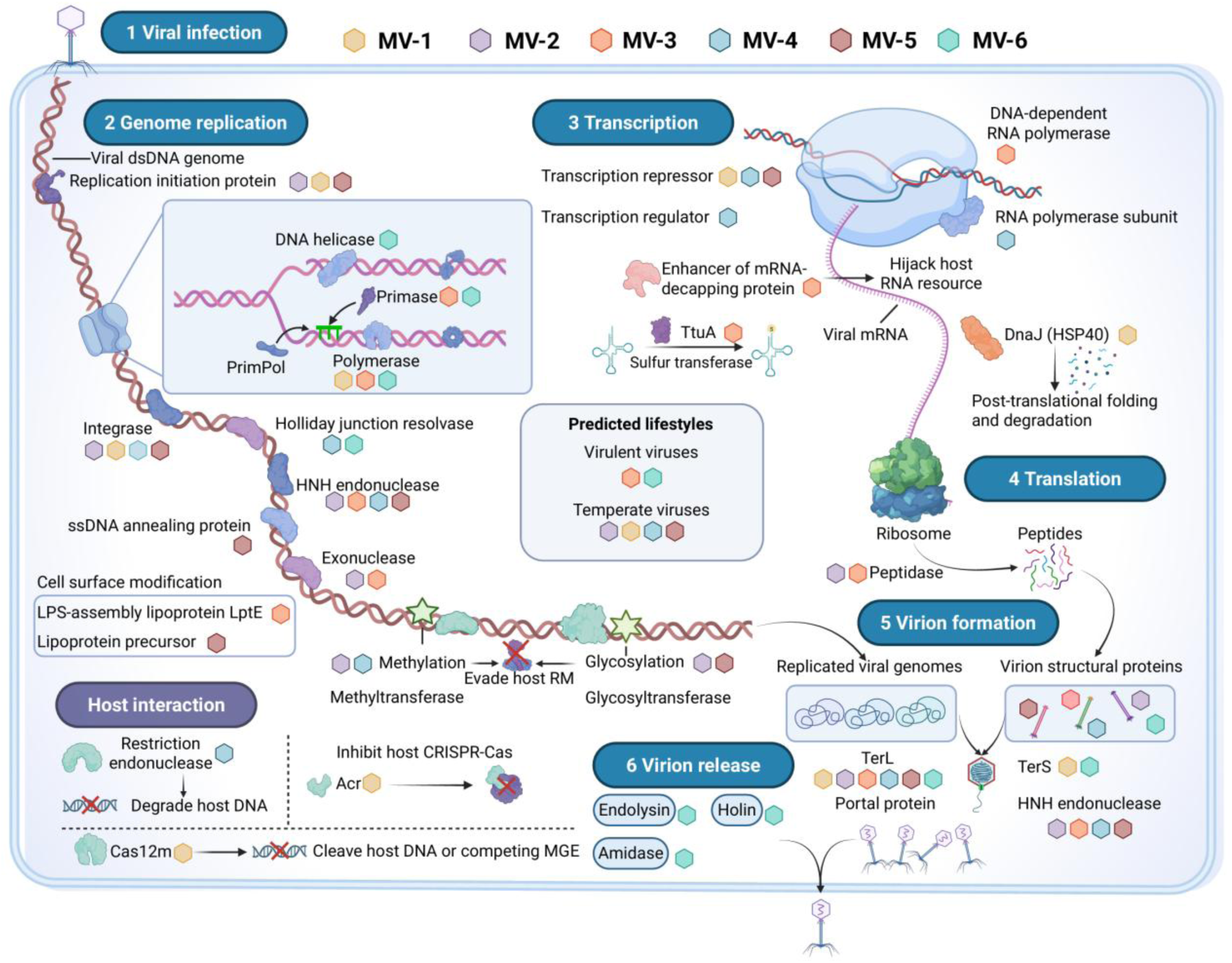
Proposed mechanisms underlying myxophage infection and interactions with myxobacterial hosts. Schematic overview of the putative mechanisms by which the six proposed myxobacterial viruses (MVs) interact with their hosts throughout infection. Colored hexagons represent different viral groups. The model summarizes the major functional modules identified in viral genomes, including genome replication, transcriptional and translational regulation, host metabolic reprogramming through auxiliary metabolic genes (AMGs), immune evasion mediated by DNA methylation and anti-CRISPR (Acr) proteins, integration and lysogeny, virion assembly, host cell lysis, and interactions with competing mobile genetic elements (MGEs). The proposed type V-M CRISPR–Cas system encoded by MV-1 is highlighted as a potential mechanism for mediating virus–host and virus–virus interactions. The diagram integrates genomic predictions and functional annotations presented in this study and is intended as a conceptual model. The illustration was created with BioRender.com.

The distinctive physiology of myxobacteria provides an ecological context for interpreting these viral adaptations. Myxobacteria rely on coordinated multicellular behaviors, cooperative predation, and extensive secondary metabolism, all of which require substantial metabolic investment. Viral genes involved in sulfur metabolism, iron homeostasis, and cell-envelope remodeling may therefore facilitate infection within this unique physiological background^109^, although their precise functions remain to be experimentally validated. Similarly, the virus-encoded CRISPR–Cas system may contribute to interactions with the host or competing mobile genetic elements, but its biological role awaits functional characterization.

In conclusion, this work substantially expands the known diversity of myxobacterial viruses and provides new insights into the molecular strategies underlying virus–host interactions in predatory bacteria. Beyond establishing a genomic framework for future studies of myxophages, these findings provide a foundation for exploring how viruses influence the ecology, evolution, and biotechnological applications of myxobacteria, including their potential use as biocontrol agents against antimicrobial-resistant pathogens.

## Methods

### Myxobacterial genome collection

Myxococcota genomes were compiled from several publicly available databases, including the NCBI GenBank database, the Genomes from Earth’s Microbiomes (GEM) catalog^110^, the Global Ocean Microbiome genome Catalogue (GOMC)^111^, the Genome Resolved Open Watersheds database (GROWdb)^112^, the Tibetan Glacier Genome and Gene (TG2G) catalog^113^, the soil microbiomes (SMAG)^114^, and the Crop Root Bacterial genome Collection (CRBC)^115^. Genomes retrieved from NCBI GenBank were selected according to the criteria outlined in the GTDB R226 database^116^: (1) taxonomically annotated as “Myxococcota”^1,117–119^. (2) quality criteria including CheckM2 completeness >50%, contamination <10%, quality score >50 (defined as completeness - 5*contamination), with >40% of bac120 marker genes, <1000 contigs, N50 >5kb, and <100,000 ambiguous bases. For other microbial genome catalogs, genomes were subjected to separate quality control using CheckM2^120^, retaining only genomes with completeness >50%, contamination <10%, and quality score >50. Taxonomic classification was performed using the classify_wf workflow in GTDB-Tk v2.4.0^121^.

### Functional annotation of myxobacterial genomes (BGCs, AMPs, DSs, and TXSSs)

Antimicrobial peptides (AMPs) were identified following a previously published workflow^122^. Brieflly, small open reading frames (smORFs) were predicted using prodigal v2.6.3^123^ (-p meta -g 11 -n -q) and filtered to retain only complete smORFs (≥33bp ≤303bp partial=00). Candidate AMPs were subsequently predicted using Macrel v1.5.0^124^ (peptides mode). Anti-phage defense systems were identified using DefenseFinder v2.0.0^125^ with MacSyFinder model v2.0.2^126^. Biosynthetic Gene Clusters (BGCs) were predicted using antiSMASH v8.0.1^127^ (--taxon bacteria --allow-long-headers). Secretion systems were identified using TXSScan v1.1.3^43,128^ with the corresponding MacSyFinder models^126^ (--db-type gembase --models TXSScan all --replicon-topology linear).

### Identification of mobile genetic elements

Plasmids and prophage-like regions were identified from Myxococcota genomes using geNomad v1.8.0^129^ (end-to-end). Regions classified as proviruses by geNomad were treated as prophage in subsequent analyses. To minimize host sequence contamination, proviral boundaries were refined using CheckV, which trims host-derived regions flanking predicted proviruses. Integrons were identified using IntegronFinder v2.0.5^130^ (--local-max --func-annot --gbk --pdf) and only complete integrons containing both an integron integrase and adjacent *attC* sites were retained. Integrative Conjugative Elements (ICEs) and Integrative Mobilizable Elements (IMEs) were identified using ICEfinder v2.0^48^. Insert sequences (ISs) were identified using ISfinder^47^, and only complete ISs were retained. Putative composite transposons were defined as pairs of IS elements separated by no more than 10 kb on the same genomic sequence.

### Identification of CRISPR arrays and CRISPR–Cas systems in myxobacterial genomes

CRISPR arrays were initially identified using MinCED^131^ and subsequently validated with CRISPRCasFinder v4.3.2^132^. Only arrays with evidence level 4 (repeat conservation index >70% and spacer identity <8%) were retained for downstream analysis. CRISPR–Cas systems located on contigs containing validated CRISPR arrays were identified using CRISPRCasTyper v1.8.0 (--prodigal meta --no_grid)^133^. To lower bias, manual curation was performed to the contigs that cannot be assigned to Myxococcota bacteria. Specifically, the predicted proteins encoded on CRISPR-containing contigs were searched against the NCBI NR database using BLASTp^134^, retaining only the best hit for each protein. Contigs were discarded when more than 50% of their predicted proteins were assigned outside the phylum Myxococcota.

### Identification and functional annotation of viruses

Myxobacterial viruses were identified from the IMG/VR v4 database^51^, the VIRE database^52^, Virus-Host DB^53^ (accessed on 8 December, 2025), myxobacterial genomes, and provirus-based methods. Validated CRISPR spacers were searched against the IMG/VR v4 and VIRE databases using BLASTn^134^ (-evalue 1e-5 -word_size 8 -task blastn-short). Viral sequences showing 100% spacer coverage with no more than one mismatch were retained, yielding 1,691 viral sequences. Previously reported myxophages were retrieved from Virus-Host DB by querying “Myxococcota”, resulting in five reference viruses (Mx8, Mx4, Mx9, Mx1, Mx4 ts27htf-1hrm-1).

Viral sequences within myxobacterial genomes were identified using geNomad v1.8.1^129^ (end-to-end), with further confirmation through CheckV v1.0.3^135^. A total 253 viral regions from bacterial genomes were retained using the same methodology applied to IMG/VR and VIRE viruses. Proviruses were identified when viral regions were flanked by two host regions and assigned high-quality status by CheckV^135^. To ensure the origination of provirus from predatory bacteria, we manually checked the taxonomy of host regions by searching host proteins against the NCBI NR database using diamond v2.1.0^136^ blastp (-k 1 -e 0.000001) with the best hit, and 51 proviruses were retained by the provirus-based method.

All 2,000 viral sequences were dereplicated using the CheckV python scripts with 95% average nucleotide identity and 85% alignment coverage^135^, resulting in 1,112 nonredundant viruses. Viral sequences were subsequently filtered using geNomad v1.8.0^129^ and CheckV v1.0.1^135^ following previously established criteria^137^. Briefly, geNomad predictions between 5 and 10 kb were required to have a virus score ≥ 0.9, at least one viral hallmark gene, and virus-marker enrichment > 2.0. Predictions ≥ 10 kb were required to have a virus score ≥ 0.8 together with at least one viral hallmark gene or virus-marker enrichment > 5.0. Contigs containing direct terminal repeats (DTRs) or inverted terminal repeats (ITRs) were also retained. All candidate viral sequences were subsequently refined with CheckV, retaining only contigs containing at least one viral gene. This workflow yielded a final dataset of 790 nonredundant myxobacterial vOTUs.

Although CRISPR spacer matches are currently the most reliable and straightforward approach to link metagenomic viral sequences to their likely hosts, there is also caveats associated with this methodology whereby non-infection interactions can also lead to CRISPR-spacer gain events^138^. To further reduce the false positives, we buttress the results with other lines of evidence using iPHoP v1.4.2^139^ (--min_score 90), which integrates multiple complementary host prediction strategies, including gene flux between viruses and the putative hosts, CRISPR spacer matches, and k-mer composition similarity. Based on iPHoP results, none of the predicted hosts belonged to taxa outside the phylum Myxococcota. Consequently, all 790 vOTUs were retained for downstream analyses.

The presence of direct terminal repeats (DTRs) or inverted terminal repeats (ITRs) was determined using geNomad^129^. Sequences containing DTRs or ITRs were considered as representing circular or linear complete genomes. Complete genomes were visualized using Proksee^140^ to provide a comprehensive view of the genomic features. Viral lifestyles (virulent/temperate) were inferred using the geNomad, CheckV and BACPHLIP v0.9.6^63^ tools. Integrated proviruses identified by both geNomad and CheckV are considered as temperate viruses. The remaining vOTUs were classified using BACPHLIP, and predictions with a greater probability (>90%) were classified as either virulent or temperate.

Predicted viral proteins were functionally annotated using HH-suite3^141^. Multiple sequence alignments were generated using HHblits v3.3.0 against the UniRef30_2023_02 database (three MSA generation iterations; e-value, 1e-6). The resulting MSAs were searched against the PDB70_May_2025^142^, PfamA 35.0^143^, NCBI CDD v3.19^144^, PHROG v4^145^, SCOPe70 v2.08^146^, and UniProt-SwissProt-viral70_Nov_2021^147^ databases using HHsearch v3.3.0 (-Z 250 -loc -z 1 -b 1 -B 250 -ssm 2 -sc 1 -seq 1 -norealign -maxres 32000).

### Viral auxiliary metabolic genes (AMGs) identification

AMGs were identified using two complementary pipelines, DRAM-v v1.5.0^148^ and VIBRANT^149^, followed by stringent genomic-context and functional filtering to generate a high-confidence AMG catalog. For the DRAM-v pipeline, viral genomes were first processed with VirSorter2 (parameter: --prep-for-dramv) to generate the required input files and then annotated using DRAM-v (parameter: --prodigal_mode meta). Candidate AMGs were curated following the criteria of Tian et al^150^. Briefly, genes assigned a DRAM-v metabolism flag (M) and flanked on both sides by viral hallmark or viral-like genes (auxiliary scores ≤ 3) were retained as candidate AMGs. Candidates were further filtered by removing AMGs derived from viral genomes harboring transposons or near the ends of viral genomes.

For VIBRANT pipeline, AMG candidates were curated according to Zhou et al^151^. Specifically, we excluded (1) genes located at contig edges; (2) genes with KEGG or Pfam v-scores ≥ 1; (3) genes whose four upstream or downstream flanking genes all had KEGG v-scores < 0.25; and (4) genes assigned to COG functional categories T or B. The remaining candidates were retained as high-confidence candidates.

AMGs identified by either pipeline and passing all filtering criteria were combined to generate the final high-confidence AMG catalog. Following the recommendations of Martin et al^152^, genes involved in DNA modification or queuosine biosynthesis, including *dcm* (DNA cytosine methyltransferase), *queC*, and *queDEF,* were excluded from AMG list. To specifically investigate viral contributions to biogeochemical cycling, the final AMG catalog was queried against CCycDB v2.1^153^ using CCycdb.pl (https://github.com/ccycdb/ccycdb.github.io), retaining only genes associated with carbon, nitrogen, sulfur, and phosphorus cycling.

### Genome-based taxonomic assignment of viruses

Gene-sharing relationships among viruses was inferred using vConTACT3 v3.1.6^154^, incorporating the viral genomes identified in this study, five previously reported myxococcota phages^27^ (Mx8, Mx4, Mx9, Mx1, Mx4 ts27htf-1hrm-1), and reference prokaryotic viruses from NCBI RefSeq (release 230). The resulting network was visualized in Cytoscape v3.10.3^155^, and genome-scale comparisons were performed using the clinker module implemented in CAGECAT v1.0^156^.

Proteome-based phylogenetic analyses were conducted using the ViPTree server^157^ with complete viral genomes belonging to the class *Caudoviricetes*. For visualization, representative subtrees containing the newly identified viruses and their closest relatives were extracted from the complete phylogeny according to the genome similarity (*SG*) metric. Candidate viral groups were considered to represent distinct viral families when they formed coherent clusters in the vConTACT3 network and independent monophyletic lineages in the ViPTree phylogeny with branch lengths exceeding the established family-level threshold (0.05)^38^. The fraction of shared orthologous proteins between viral genomes was estimated using CompareM (https://github.com/dparks1134/CompareM) (-evalue 1e-5 -identity 30%). Intergenomic nucleotide similarity between newly identified complete genomes and reference viruses was calculated using VIRIDIC^59^, which implements the traditional algorithm used by the International Committee on Taxonomy of Viruses (ICTV) to calculate virus intergenomic similarities.

### Protein structural prediction and network analysis

Protein structures pf viral Cas12m, and Acr proteins were predicted using the AlphaFold3 web server^158^. Major capsid proteins, portal proteins, and terminase large subunits sequence similarity networks were generated using CLANS (BLASTp option -evalue 1e-4)^159^ and visualized with Cytoscape^155^.

### Identification of viral CRISPR–Cas systems

CRISPR–Cas systems encoded by viral genomes were identified using CRISPRCasTyper^133^. Viral CRISPR spacers were searched against the IMG/VR v4 database, the IMG/PR database, and the myxobacterial genome collection using BLASTn, but no significant matches were detected. To examine catalytic residue conservation, the Cas12m protein encoded by MV-1 was aligned with previously characterized bacterial Cas12m proteins^160^ using Clustal Omega^161^, and the alignment was visualized with ESPript 3.0^162^. The three-dimensional structure of MV-1 Cas12m was predicted using AlphaFold3^158^ and visualized with ChimeraX^163^.

### Phylogenetic analysis

Phylogenomic relationships of myxobacterial genomes encoding CRISPR–Cas systems or predicted to serve as viral hosts were reconstructed using three Cyanobacteriota genomes as outgroups. The 120 concatenated bacterial marker genes identified by the GTDB-Tk classify_wf module^121^ were concatenated and used to infer a maximum-likelihood phylogeny with the GTDB-Tk^121^ infer workflow. The resulting tree was rooted using the Cyanobacteriota outgroup and visualized in iTOL^164^.

For Cas12m evolution, the MV-1 Cas12m were aligned with representative type V CRISPR effectors from archaea^165^, bacteria^107,160^, bacteriophage^95^ and bacterial TnpB nucleases (derived from NCBI) using MUSCLE v5.3^166^. The resulting multiple sequence alignments were trimmed with trimAl v1.5^167^ (-gt 0.5), and a maximum likelihood phylogeny was reconstructed using iqtree v2.4.0^168^ (-m MFP -bb 1000 -nt AUTO). The tree was rooted with the TnpB clade and visualized with iTOL^164^.

## Data Availability

All data supporting this study are available. Genomes of myxobacteria are accessible from the NCBI GenBank database (https://www.ncbi.nlm.nih.gov/genbank/), the Genomes from Earth’s Microbiomes catalogue (https://portal.nersc.gov/GEM/), the Global Ocean Microbiome genome Catalogue (https://db.cngb.org/maya/datasets/MDB0000002), the Genome Resolved Open Watersheds database (https://doi.org/10.5281/zenodo.11193259), the Tibetan Glacier Genome and Gene catalog (https://www.biosino.org/node/project/detail/OEP00003083), the soil microbiomes (https://doi.org/10.5281/zenodo.8223844), and the Crop Root Bacterial genome Collection (https://www.cropmicrobiome.com/). CRISPR arrays, as well as genome sequences of viruses identified in this study have been deposited in Zenodo (https://doi.org/10.5281/zenodo.17373696). Relevant sample attributes (e.g., sampling locations and ecosystem types) were obtained from IMG/VR or NCBI BioSample (https://www.ncbi.nlm.nih.gov/biosample/). All databases used in this study (UniRef30, PfamA 35.0, NCBI CDD v3.19, PDB70_May_2025, PHROG v4, SCOPe70 v2.08 and UniProt-SwissProt-viral70_Nov_2021) are publicly available at https://wwwuser.gwdguser.de/~compbiol/data/hhsuite/databases/hhsuite_dbs/.

## Code Availability

Wrapper scripts supporting all key analyses of this work are available on GitHub (https://github.com/lianmsu/Predatory-bacterial-virome).

## Acknowledgements

This work was supported by the National Natural Science Foundation of China (425B2048, 51721006, and 92047303). Supports from the High-performance Computing Platform of Peking University are acknowledged.

## Author Contributions

J.R.N. designed the research. P.W.L. conducted the bioinformatic analysis. P.W.L. wrote the manuscript and J.R.N. revised the manuscript. All the authors read and approved the final manuscript.

## Competing Interests

The authors declare no competing interests.

## Notes

### Competing Interest Statement

The authors have declared no competing interest.

## References

1. Zhang, L., Guo, L., Cui, Z. & Ju, F. Exploiting predatory bacteria as biocontrol agents across ecosystems. Trends Microbiol. 32, 398–409 (2024).

2. Vasse, M. & Velicer, G. J. Predation in microbial communities: gradients of nutritive killing. Nat. Rev. Microbiol. 1–20 (2026) doi:10.1038/s41579-026-01299-7.

3. Pérez, J., Moraleda-Muñoz, A., Marcos-Torres, F. J. & Muñoz-Dorado, J. Bacterial predation: 75 years and counting! *Environ*. Microbiol. 18, 766–779 (2016).

4. Vasse, M., Fiegna, F., Kriesel, B. & Velicer, G. J. Killer prey: ecology reverses bacterial predation. PLOS Biol. 22, e3002454 (2024).

5. Ledvina, H. E. et al. Functional amyloid proteins confer defence against predatory bacteria. Nature 644, 1–8 (2025).

6. Hofer, U. Bacteria on the hunt. Nat. Rev. Microbiol. 19, 406–406 (2021).

7. Hungate, B. A. et al. The Functional Significance of Bacterial Predators. mBio 12, e00466–21 (2021).

8. Atterbury, R. J. & Tyson, J. Predatory bacteria as living antibiotics – where are we now? Microbiology 167, (2021).

9. Sockett, R. E. Learning with bdellovibrio. Nat. Microbiol. 8, 1189–1190 (2023).

10. Caulton, S. G. et al. Bdellovibrio bacteriovorus uses chimeric fibre proteins to recognize and invade a broad range of bacterial hosts. Nat. Microbiol. 9, 214–227 (2024).

11. Zhang, L., Huang, X., Zhou, J. & Ju, F. Active predation, phylogenetic diversity, and global prevalence of myxobacteria in wastewater treatment plants. ISME J. 17, 671–681 (2023).

12. Muñoz-Dorado, J., Marcos-Torres, F. J., García-Bravo, E., Moraleda-Muñoz, A. & Pérez, J. Myxobacteria: moving, killing, feeding, and surviving together. Front. Microbiol. 7, 781 (2016).

13. Alexakis, K., Baliou, S. & Ioannou, P. Predatory bacteria in the treatment of infectious diseases and beyond. Infect. Dis. Rep. 16, 684–698 (2024).

14. Whitworth, D. E., Sydney, N. & Radford, E. J. Myxobacterial Genomics and Post-Genomics: A Review of Genome Biology, Genome Sequences and Related ‘Omics Studies. Microorganisms 9, 2143 (2021).

15. Berleman, J. E. & Kirby, J. R. Deciphering the hunting strategy of a bacterial wolfpack. FEMS Microbiol. Rev. 33, 942–957 (2009).

16. Salamzade, R. A., Kalan, L. R. & Currie, C. R. Complex multicellularity is linked with expanded specialized metabolite production in microorganisms. Nat. Microbiol. 1–13 (2026) doi:10.1038/s41564-026-02385-5.

17. Schneiker, S. et al. Complete genome sequence of the myxobacterium sorangium cellulosum. Nat. Biotechnol. 25, 1281–1289 (2007).

18. Villegas, C., et al. Epothilones as natural compounds for novel anticancer drugs development. Int. J. Mol. Sci. 24, 6063 (2023).

19. Sharma, G., Khatri, I. & Subramanian, S. Complete genome of the starch-degrading myxobacteria sandaracinus amylolyticus DSM 53668T. Genome Biol. Evol. 8, 2520–2529 (2016).

20. Mohr, K. I. Diversity of myxobacteria—we only see the tip of the iceberg. Microorganisms 6, 84 (2018).

21. Weissman, K. J. & Müller, R. Myxobacterial secondary metabolites: bioactivities and modes-of-action. Nat. Prod. Rep. 27, 1276–1295 (2010).

22. Södergren, J. et al. Myxobacteria isolated from recirculating aquaculture systems (RAS): ecology and significance as off-flavor producers. Appl. Environ. Microbiol. 91, e00757–25 (2025).

23. Swetha, R. G. et al. MyxoPortal: a database of myxobacterial genomic features. Database 2024, baae056 (2024).

24. Dai, W., Liu, Y., Cui, Z., Li, W. & Wang, H. From predation to function: how myxobacteria drive soil microbial community dynamics and ecological functions. Appl. Environ. Microbiol. 91, e01922–25 (2025).

25. Chevallereau, A., Pons, B. J., Van Houte, S. & Westra, E. R. Interactions between bacterial and phage communities in natural environments. Nat. Rev. Microbiol. 20, 49–62 (2022).

26. Roux, S. & Coclet, C. Viromics approaches for the study of viral diversity and ecology in microbiomes. Nat. Rev. Genet. 1–15 (2025) doi:10.1038/s41576-025-00871-w.

27. López-Rojo, A. et al. Genome sequences of Mx1, the first myxococcus phage isolated, and Mx4, a generalized transducing myxophage. Microbiol. Resour. Announce. 12, e00904–23 (2023).

28. Vasse, M. & Wielgoss, S. Bacteriophages of myxococcus xanthus, a social bacterium. Viruses 10, 374 (2018).

29. Martin, S., Sodergren, E., Masuda, T. & Kaiser, D. Systematic isolation of transducing phages for myxococcus xanthus. Virology 88, 44–53 (1978).

30. Wang, J. Y. & Doudna, J. A. CRISPR technology: a decade of genome editing is only the beginning. Science 379, eadd8643 (2023).

31. Duan, C. et al. Diversity of bathyarchaeia viruses in metagenomes and virus-encoded CRISPR system components. ISME Commun. 4, ycad011 (2024).

32. Zhou, Y. et al. Viruses and virus satellites of haloarchaea and their nanosized DPANN symbionts reveal intricate nested interactions. Nat. Microbiol. 1–13 (2025) doi:10.1038/s41564-025-02149-7.

33. Liu, J., Jaffe, A. L., Chen, L., Bor, B. & Banfield, J. F. Host translation machinery is not a barrier to phages that interact with both CPR and non-CPR bacteria. mBio 14, e01766–23 (2023).

34. Medvedeva, S. et al. Three families of asgard archaeal viruses identified in metagenome-assembled genomes. Nat. Microbiol. 7, 962–973 (2022).

35. Rambo, I. M., Langwig, M. V., Leão, P., De Anda, V. & Baker, B. J. Genomes of six viruses that infect asgard archaea from deep-sea sediments. Nat. Microbiol. 7, 953–961 (2022).

36. Wu, Z., Liu, S. & Ni, J. Metagenomic characterization of viruses and mobile genetic elements associated with the DPANN archaeal superphylum. Nat. Microbiol. 9, 3362–3375 (2024).

37. Laso-Pérez, R. et al. Evolutionary diversification of methanotrophic ANME-1 archaea and their expansive virome. Nat. Microbiol. 8, 231–245 (2023).

38. Medvedeva, S., Borrel, G., Krupovic, M. & Gribaldo, S. A compendium of viruses from methanogenic archaea reveals their diversity and adaptations to the gut environment. Nat. Microbiol. 8, 2170–2182 (2023).

39. Zuckerman, D. M., So, J. M. T. & Hoiczyk, E. Secretins of type-two secretion systems are necessary for exopolymeric slime secretion in cyanobacteria and myxobacteria. Nat. Commun. 16, 8482 (2025).

40. Georjon, H. & Bernheim, A. The highly diverse antiphage defence systems of bacteria. Nat. Rev. Microbiol. 21, 686–700 (2023).

41. Li, P. et al. The defensome of prokaryotes in aquifers. Nat. Commun. 16, 6482 (2025).

42. Treuner-Lange, A., et al. Tight-packing of large pilin subunits provides distinct structural and mechanical properties for the myxococcus xanthus type IVa pilus. Proc. Natl. Acad. Sci. 121, e2321989121 (2024).

43. Denise, R., Abby, S. S. & Rocha, E. P. C. Diversification of the type IV filament superfamily into machines for adhesion, protein secretion, DNA uptake, and motility. PLOS Biol. 17, e3000390 (2019).

44. Arnold, B. J., Huang, I.-T. & Hanage, W. P. Horizontal gene transfer and adaptive evolution in bacteria. Nat. Rev. Microbiol. 20, 206–218 (2022).

45. Guo, J. et al. Mobile genetic elements shape microbial diversity and functions in thawing permafrost soils. Nat. Microbiol. 11, 1800–1814 (2026).

46. Lang, A. S., Buchan, A. & Burrus, V. Interactions and evolutionary relationships among bacterial mobile genetic elements. Nat. Rev. Microbiol. 23, 423–438 (2025).

47. Siguier, P., Perochon, J., Lestrade, L., Mahillon, J. & Chandler, M. ISfinder: the reference centre for bacterial insertion sequences. Nucleic Acids Res. 34, D32–36 (2006).

48. Wang, M. et al. ICEberg 3.0: functional categorization and analysis of the integrative and conjugative elements in bacteria. Nucleic Acids Res. 52, D732–D737 (2024).

49. Harrison, L. et al. Pangenomic characterization of campylobacter plasmids for enhanced molecular typing, risk assessment and source attribution. Pathogens 14, 936 (2025).

50. Hu, W. et al. Characteristics and immune functions of the endogenous CRISPR-cas systems in myxobacteria. mSystems 9, e01210–23 (2024).

51. Camargo, A. P. et al. IMG/VR v4: an expanded database of uncultivated virus genomes within a framework of extensive functional, taxonomic, and ecological metadata. Nucleic Acids Res. 51, D733–D743 (2023).

52. Nishijima, S., Fullam, A., Schmidt, T. S. B., Kuhn, M. & Bork, P. VIRE: a metagenome-derived, planetary-scale virome resource with environmental context. Nucleic Acids Res. gkaf1225 (2025) doi:10.1093/nar/gkaf1225.

53. Mihara, T. et al. Linking virus genomes with host taxonomy. Viruses 8, 0–6 (2016).

54. Wielgoss, S. & Julien, B. Whole-genome sequence of *Myxococcus* phage Mx9. Microbiol. Resour. Announce. 12, e00221–23 (2023).

55. Wang, Y. et al. Early transcriptome of pseudomonas aeruginosa PAO1 infected with vB_Pae_QDWS, a short-latent lytic phage. J. Basic Microbiol. 65, e70105 (2025).

56. Cumby, N., Davidson, A. R. & Maxwell, K. L. The moron comes of age. Bacteriophage 2, 225–228 (2012).

57. Lu, A. et al. Exopolysaccharide biosynthesis genes required for social motility in myxococcus xanthus. Mol. Microbiol. 55, 206–220 (2005).

58. Ackermann, H.-W., Krisch, H. M. & Comeau, A. M. Morphology and genome sequence of phage φ1402. Bacteriophage 1, 138–142 (2011).

59. Moraru, C., Varsani, A. & Kropinski, A. M. VIRIDIC—a novel tool to calculate the intergenomic similarities of prokaryote-infecting viruses. Viruses 12, (2020).

60. Martins, S. J. et al. Predators of soil bacteria in plant and human health. Phytobiomes J. 6, 184–200 (2022).

61. Shapiro, O. H., Kushmaro, A. & Brenner, A. Bacteriophage predation regulates microbial abundance and diversity in a full-scale bioreactor treating industrial wastewater. ISME J. 4, 327–336 (2010).

62. Mohsenipour, Z. et al. Predation on bacterial pathogens by predatory bacteria of sewage origin: three days prey-predator interactions. BMC Microbiol. 24, 516 (2024).

63. Hockenberry, A. J. & Wilke, C. O. BACPHLIP: predicting bacteriophage lifestyle from conserved protein domains. PeerJ 9, e11396 (2021).

64. Groth, A. C. & Calos, M. P. Phage integrases: biology and applications. J. Mol. Biol. 335, 667–678 (2004).

65. Lewis, D., Le, P., Zurla, C., Finzi, L. & Adhya, S. Multilevel autoregulation of λ repressor protein CI by DNA looping in vitro. Proc. Natl. Acad. Sci. 108, 14807–14812 (2011).

66. Bennett, R. J., Dunderdale, H. J. & West, S. C. Resolution of holliday junctions by RuvC resolvase: cleavage specificity and DNA distortion. Cell 74, 1021–1031 (1993).

67. Kuzminov, A. Recombinational repair of DNA damage in escherichia coli and bacteriophage lambda. Microbiol. Mol. Biol. Rev.: MMBR 63, 751–813, table of contents (1999).

68. Kala, S. et al. HNH proteins are a widespread component of phage DNA packaging machines. Proc. Natl. Acad. Sci. 111, 6022–6027 (2014).

69. Kazlauskas, D., Krupovic, M., Guglielmini, J., Forterre, P. & Venclovas, Č. Diversity and evolution of B-family DNA polymerases. Nucleic Acids Research 48, 10142–10156 (2020).

70. Kazlauskas, D., Krupovic, M. & Venclovas, Č. The logic of DNA replication in double-stranded DNA viruses: insights from global analysis of viral genomes. Nucleic Acids Res. 44, 4551–4564 (2016).

71. Guilliam, T. A., Keen, B. A., Brissett, N. C. & Doherty, A. J. Primase-polymerases are a functionally diverse superfamily of replication and repair enzymes. Nucleic Acids Res 43, 6651–6664 (2015).

72. Valero-Rello, A. Diversity, specificity and molecular evolution of the lytic arsenal of pseudomonas phages: in silico perspective. Environ. Microbiol. 21, 4136–4150 (2019).

73. Urvoy, M., Baumgart, L., Howard-Varona, C. & Sullivan, M. B. Beyond AMGs: phage-encoded transcription and sigma factors as understudied virocell reprogramming tools. Trends Microbiol. 0, (2026).

74. Cantu, F. et al. Poxvirus-encoded decapping enzymes promote selective translation of viral mRNAs. PLOS Pathog. 16, e1008926 (2020).

75. Tran Thi Ngoc, A., Nguyen Van, K. & Lee, Y. H. DnaJ, a heat shock protein 40 family member, is essential for the survival and virulence of plant pathogenic *pseudomonas cichorii* JBC1. Res. Microbiol. 174, 104094 (2023).

76. Knox, C., Luke, G. A., Blatch, G. L. & Pesce, E.-R. Heat shock protein 40 (Hsp40) plays a key role in the virus life cycle. Virus Res. 160, 15–24 (2011).

77. Chen, M. et al. The [4Fe-4S] cluster of sulfurtransferase TtuA desulfurizes TtuB during tRNA modification in thermus thermophilus. *Commun*. Biol. 3, 168 (2020).

78. Huang, D. et al. Adaptive strategies and ecological roles of phages in habitats under physicochemical stress. Trends Microbiol. 32, 902–916 (2024).

79. Lee, K., Bekiari, D., Doello, S. & Forchhammer, K. The (glg)ABCs of cyanobacteria: modelling of glycogen synthesis and functional divergence of glycogen synthases in synechocystis sp. PCC 6803. FEBS Lett. 1–18 doi:10.1002/1873-3468.70299.

80. Yuan, L. & Ju, F. Potential auxiliary metabolic capabilities and activities reveal biochemical impacts of viruses in municipal wastewater treatment plants. Environ. Sci. Technol. 57, 5485–5498 (2023).

81. Spigelmyer, S. M. & Santos, P. C. D. Intricacies in iron–sulfur cluster function and biogenesis: functional versatility, sulfur sources, and enzyme specificity. *RSC Chem*. Biol. 7, 763–782 (2026).

82. Zhao, X., Zhou, Y., Zhang, Y. & Zhang, Y. Ferritin: significance in viral infections. Rev. Med. Virol. 34, e2531 (2024).

83. Songire, V. M. & Patil, R. H. Microbial antioxidative enzymes: biotechnological production and environmental and biomedical applications. Appl. Biochem. Microbiol. 61, 1– 26 (2025).

84. Whitworth, D. E. Myxobacteria: physiology and regulation. Microorganisms 10, 805 (2022).

85. Bertozzi Silva, J., Storms, Z. & Sauvageau, D. Host receptors for bacteriophage adsorption. FEMS Microbiol. Lett. 363, fnw002 (2016).

86. Rousset, F. et al. Phages and their satellites encode hotspots of antiviral systems. Cell Host Microbe 30, 741–753.e5 (2022).

87. Pathak, D. T., Wei, X., Dey, A. & Wall, D. Molecular recognition by a polymorphic cell surface receptor governs cooperative behaviors in bacteria. PLOS Genet. 9, e1003891 (2013).

88. Jeudy, S. et al. The DNA methylation landscape of giant viruses. Nat. Commun. 11, 2657 (2020).

89. Murphy, J., Mahony, J., Ainsworth, S., Nauta, A. & van Sinderen, D. Bacteriophage orphan DNA methyltransferases: insights from their bacterial origin, function, and occurrence. Appl. Environ. Microbiol. 79, 7547–7555 (2013).

90. Markine-Goriaynoff, N. et al. Glycosyltransferases encoded by viruses. J. Gen. Virol. 85, 2741–2754 (2004).

91. Beavogui, A. et al. The defensome of complex bacterial communities. Nat. Commun. 15, 2146 (2024).

92. Makarova, K. S. et al. Evolutionary classification of CRISPR–cas systems: a burst of class 2 and derived variants. Nat. Rev. Microbiol. 18, 67–83 (2020).

93. Wu, W. Y., Adiego-Pérez, B. & van der Oost, J. Biology and applications of CRISPR– Cas12 and transposon-associated homologs. Nat. Biotechnol. 42, 1807–1821 (2024).

94. Wu, W. Y. et al. The miniature CRISPR-Cas12m effector binds DNA to block transcription. Mol. Cell 82, 4487–4502.e7 (2022).

95. Pausch, P., et al. CRISPR-CasΦ from huge phages is a hypercompact genome editor. Science 369, 333–337 (2020).

96. Altae-Tran, H. et al. Diversity, evolution, and classification of the RNA-guided nucleases TnpB and Cas12. Proc. Natl. Acad. Sci. 120, e2308224120 (2023).

97. Faure, G. et al. CRISPR–cas in mobile genetic elements: counter-defence and beyond. Nat. Rev. Microbiol. 17, 513–525 (2019).

98. Medvedeva, S. et al. Virus-borne mini-CRISPR arrays are involved in interviral conflicts. Nat. Commun. 10, 5204 (2019).

99. Jin, S. et al. Functional RNA splitting drove the evolutionary emergence of type V CRISPR-cas systems from transposons. Cell 0, S0092867425010359 (2025).

100. Al-Shayeb, B., et al. Diverse virus-encoded CRISPR-cas systems include streamlined genome editors. Cell 185, 4574–4586.e16 (2022).

101. Al-Shayeb, B. et al. Clades of huge phages from across earth’s ecosystems. Nature 578, 425–431 (2020).

102. Feiner, R. et al. A new perspective on lysogeny: prophages as active regulatory switches of bacteria. Nat. Rev. Microbiol. 13, 641–650 (2015).

103. Mayo-Muñoz, D., Pinilla-Redondo, R., Camara-Wilpert, S., Birkholz, N. & Fineran, P. C. Inhibitors of bacterial immune systems: discovery, mechanisms and applications. Nat. Rev. Genet. 25, 237–254 (2024).

104. Johannesman, A., Awasthi, L. C., Carlson, N. & LeRoux, M. Phages carry orphan antitoxin-like enzymes to neutralize the DarTG1 toxin-antitoxin defense system. Nat. Commun. 16, 1598 (2025).

105. Stokar-Avihail, A. et al. Discovery of phage determinants that confer sensitivity to bacterial immune systems. Cell 186, 1863–1876.e16 (2023).

106. Abergel, C., Rudinger-Thirion, J., Giegé, R. & Claverie, J.-M. Virus-encoded aminoacyl-tRNA synthetases: structural and functional characterization of mimivirus TyrRS and MetRS. J. Virol. 81, 12406–12417 (2007).

107. Yan, W. X. et al. Functionally diverse type V CRISPR-cas systems. Science 363, 88– 91 (2019).

108. Dion, M. B., Oechslin, F. & Moineau, S. Phage diversity, genomics and phylogeny. Nat. Rev. Microbiol. 18, 125–138 (2020).

109. Sargen, M. R. & Helaine, S. A prophage intercepts pathogenic activity of infecting phage for defense. Cell Host Microbe 33, 1657–1666.e4 (2025).

110. Nayfach, S. et al. A genomic catalog of Earth’s microbiomes. Nat. Biotechnol. 39, 499– 509 (2021).

111. Chen, J. et al. Global marine microbial diversity and its potential in bioprospecting. Nature 633, 371–379 (2024).

112. Borton, M. A. et al. A functional microbiome catalogue crowdsourced from north american rivers. Nature 637, 103–112 (2025).

113. Liu, Y. et al. A genome and gene catalog of glacier microbiomes. Nat Biotechnol 40, 1341–1348 (2022).

114. Ma, B. et al. A genomic catalogue of soil microbiomes boosts mining of biodiversity and genetic resources. Nat. Commun. 14, 7318 (2023).

115. Dai, R. et al. Crop root bacterial and viral genomes reveal unexplored species and microbiome patterns. Cell 188, 2521–2539.e22 (2025).

116. Parks, D. H. et al. A complete domain-to-species taxonomy for bacteria and archaea. Nat. Biotechnol. 38, 1079–1086 (2020).

117. Moreira, D., Zivanovic, Y., López-Archilla, A. I., Iniesto, M. & López-García, P. Reductive evolution and unique predatory mode in the CPR bacterium vampirococcus lugosii. Nat. Commun. 12, 2454 (2021).

118. Steichen, S. A. & Brown, J. K. Real-time quantitative detection of vampirovibrio chlorellavorus, an obligate bacterial pathogen of chlorella sorokiniana. J. Appl. Phycol. 31, 1117–1129 (2019).

119. Wang, Z., Kadouri, D. E. & Wu, M. Genomic insights into an obligate epibiotic bacterial predator: micavibrio aeruginosavorus ARL-13. BMC Genomics 12, 453 (2011).

120. Chklovski, A., Parks, D. H., Woodcroft, B. J. & Tyson, G. W. CheckM2: a rapid, scalable and accurate tool for assessing microbial genome quality using machine learning. Nat. Methods 20, 1203–1212 (2023).

121. Chaumeil, P.-A., Mussig, A. J., Hugenholtz, P. & Parks, D. H. GTDB-tk v2: memory friendly classification with the genome taxonomy database. Bioinformatics 38, 5315–5316 (2022).

122. Santos-Júnior, C. D. et al. Discovery of antimicrobial peptides in the global microbiome with machine learning. Cell 187, 3761–3778.e16 (2024).

123. Hyatt, D., LoCascio, P. F., Hauser, L. J. & Uberbacher, E. C. Gene and translation initiation site prediction in metagenomic sequences. Bioinformatics 28, 2223–2230 (2012).

124. Santos-Júnior, C. D., Pan, S., Zhao, X.-M. & Coelho, L. P. Macrel: antimicrobial peptide screening in genomes and metagenomes. PeerJ 8, e10555 (2020).

125. Tesson, F. et al. Systematic and quantitative view of the antiviral arsenal of prokaryotes. Nat. Commun. 13, 2561 (2022).

126. Néron, B. et al. MacSyFinder v2: improved modelling and search engine to identify molecular systems in genomes. Peer Community J. 3, (2023).

127. Blin, K. et al. antiSMASH 8.0: extended gene cluster detection capabilities and analyses of chemistry, enzymology, and regulation. Nucleic Acids Res. 53, gkaf334 (2025).

128. Abby, S. S. et al. Identification of protein secretion systems in bacterial genomes. Sci. Rep. 6, 23080 (2016).

129. Camargo, A. P. et al. Identification of mobile genetic elements with geNomad. Nat. Biotechnol. 42, 1303–1312 (2024).

130. Néron, B. et al. IntegronFinder 2.0: Identification and Analysis of Integrons across Bacteria, with a Focus on Antibiotic Resistance in Klebsiella. Microorganisms 10, 700 (2022).

131. Bland, C. et al. CRISPR recognition tool (CRT): a tool for automatic detection of clustered regularly interspaced palindromic repeats. BMC Bioinf. 8, 209 (2007).

132. Couvin, D. et al. CRISPRCasFinder, an update of CRISRFinder, includes a portable version, enhanced performance and integrates search for cas proteins. Nucleic Acids Res. 46, W246–W251 (2018).

133. Russel, J., Pinilla-Redondo, R., Mayo-Muñoz, D., Shah, S. A. & Sørensen, S. J. CRISPRCasTyper: automated identification, annotation, and classification of CRISPR-cas loci. CRISPR J. 3, 462–469 (2020).

134. Camacho, C. et al. BLAST+: architecture and applications. BMC Bioinf. 10, 421 (2009).

135. Nayfach, S. et al. CheckV assesses the quality and completeness of metagenome-assembled viral genomes. Nat. Biotechnol. 39, 578–585 (2021).

136. Buchfink, B., Reuter, K. & Drost, H.-G. Sensitive protein alignments at tree-of-life scale using DIAMOND. Nat. Methods 18, 366–368 (2021).

137. Graham, E. B. et al. A global atlas of soil viruses reveals unexplored biodiversity and potential biogeochemical impacts. Nat. Microbiol. 9, 1873–1883 (2024).

138. Hwang, Y., Roux, S., Coclet, C., Krause, S. J. E. & Girguis, P. R. Viruses interact with hosts that span distantly related microbial domains in dense hydrothermal mats. Nat. Microbiol. 8, 946–957 (2023).

139. Roux, S. et al. iPHoP: an integrated machine learning framework to maximize host prediction for metagenome-derived viruses of archaea and bacteria. PLOS Biol. 21, e3002083 (2023).

140. Grant, J. R. et al. Proksee: in-depth characterization and visualization of bacterial genomes. Nucleic Acids Res. 51, W484–W492 (2023).

141. Steinegger, M. et al. HH-suite3 for fast remote homology detection and deep protein annotation. BMC Bioinf. 20, 473 (2019).

142. Burley, S. K. et al. Updated resources for exploring experimentally-determined PDB structures and computed structure models at the RCSB protein data bank. Nucleic Acids Res. 53, D564–D574 (2025).

143. Finn, R. D. et al. Pfam: the protein families database. Nucleic Acids Res. 42, D222–230 (2014).

144. Marchler-Bauer, A. et al. CDD: NCBI’s conserved domain database. Nucleic Acids Res. 43, D222–D226 (2015).

145. Terzian, P., et al. PHROG: families of prokaryotic virus proteins clustered using remote homology. NAR Genomics Bioinf. 3, lqab067 (2021).

146. Chandonia, J.-M. et al. SCOPe: improvements to the structural classification of proteins - extended database to facilitate variant interpretation and machine learning. Nucleic Acids Res. 50, D553–D559 (2022).

147. UniProt Consortium. UniProt: the universal protein knowledgebase in 2023. Nucleic Acids Res. 51, D523–D531 (2023).

148. Shaffer, M. et al. DRAM for distilling microbial metabolism to automate the curation of microbiome function. Nucleic Acids Res 48, 8883–8900 (2020).

149. Kieft, K., Zhou, Z. & Anantharaman, K. VIBRANT: automated recovery, annotation and curation of microbial viruses, and evaluation of viral community function from genomic sequences. Microbiome 8, 90 (2020).

150. Tian, F. et al. Prokaryotic-virus-encoded auxiliary metabolic genes throughout the global oceans. Microbiome 12, 159 (2024).

151. Zhou, Z. et al. Unravelling viral ecology and evolution over 20 years in a freshwater lake. Nat. Microbiol. 10, 231–245 (2025).

152. Martin, C., Emerson, J. B., Roux, S. & Anantharaman, K. A call for caution in the biological interpretation of viral auxiliary metabolic genes. Nat Microbiol 10, 2122–2129 (2025).

153. Zhou, J. et al. CCycDB: an integrative knowledgebase to fingerprint microbially mediated carbon cycling processes. 2026.01.28.702190 Preprint at 10.64898/2026.01.28.702190 (2026).

154. Bolduc, B. et al. Machine learning enables scalable and systematic hierarchical virus taxonomy. Nat. Biotechnol. 1–10 (2025) doi:10.1038/s41587-025-02946-9.

155. Shannon, P. et al. Cytoscape: a software environment for integrated models of biomolecular interaction networks. Genome Res. 13, 2498–2504 (2003).

156. Gilchrist, C. L. M. & Chooi, Y.-H. clinker & clustermap.js: automatic generation of gene cluster comparison figures. Bioinformatics 37, 2473–2475 (2021).

157. Nishimura, Y. et al. ViPTree: the viral proteomic tree server. Bioinform. (oxf. Engl.) 33, 2379–2380 (2017).

158. Abramson, J. et al. Accurate structure prediction of biomolecular interactions with AlphaFold 3. Nature 630, 493–500 (2024).

159. Frickey, T. & Lupas, A. CLANS: a java application for visualizing protein families based on pairwise similarity. Bioinformatics 20, 3702–3704 (2004).

160. Bigelyte, G. et al. Innate programmable DNA binding by CRISPR-Cas12m effectors enable efficient base editing. Nucleic Acids Res. 52, 3234–3248 (2024).

161. Madeira, F. et al. Search and sequence analysis tools services from EMBL-EBI in 2022. Nucleic Acids Res. 50, W276–W279 (2022).

162. Robert, X. & Gouet, P. Deciphering key features in protein structures with the new ENDscript server. Nucleic Acids Res. 42, W320–324 (2014).

163. Meng, E. C. et al. UCSF ChimeraX: tools for structure building and analysis. Protein Sci. 32, e4792 (2023).

164. Letunic, I. & Bork, P. Interactive tree of life (iTOL) v6: recent updates to the phylogenetic tree display and annotation tool. Nucleic Acids Res. 52, W78–W82 (2024).

165. Harrington, L. B. et al. Programmed DNA destruction by miniature CRISPR-Cas14 enzymes. Science 362, 839–842 (2018).

166. Edgar, R. C. Muscle5: high-accuracy alignment ensembles enable unbiased assessments of sequence homology and phylogeny. Nat. Commun. 13, 6968 (2022).

167. Capella-Gutiérrez, S., Silla-Martínez, J. M. & Gabaldón, T. trimAl: a tool for automated alignment trimming in large-scale phylogenetic analyses. Bioinformatics 25, 1972– 1973 (2009).

168. Minh, B. Q. et al. IQ-TREE 2: new models and efficient methods for phylogenetic inference in the genomic era. Mol. Biol. Evol. 37, 1530–1534 (2020).

