## Supplementary Materials for "CRISPR mapping unveils global prevalence of viral predators of Myxobacteria"

Pengwei Li, Jinren Ni*

**This PDF file includes:**

Figs. S1 to S8
Tables S1 to S14

### Supplementary Figures

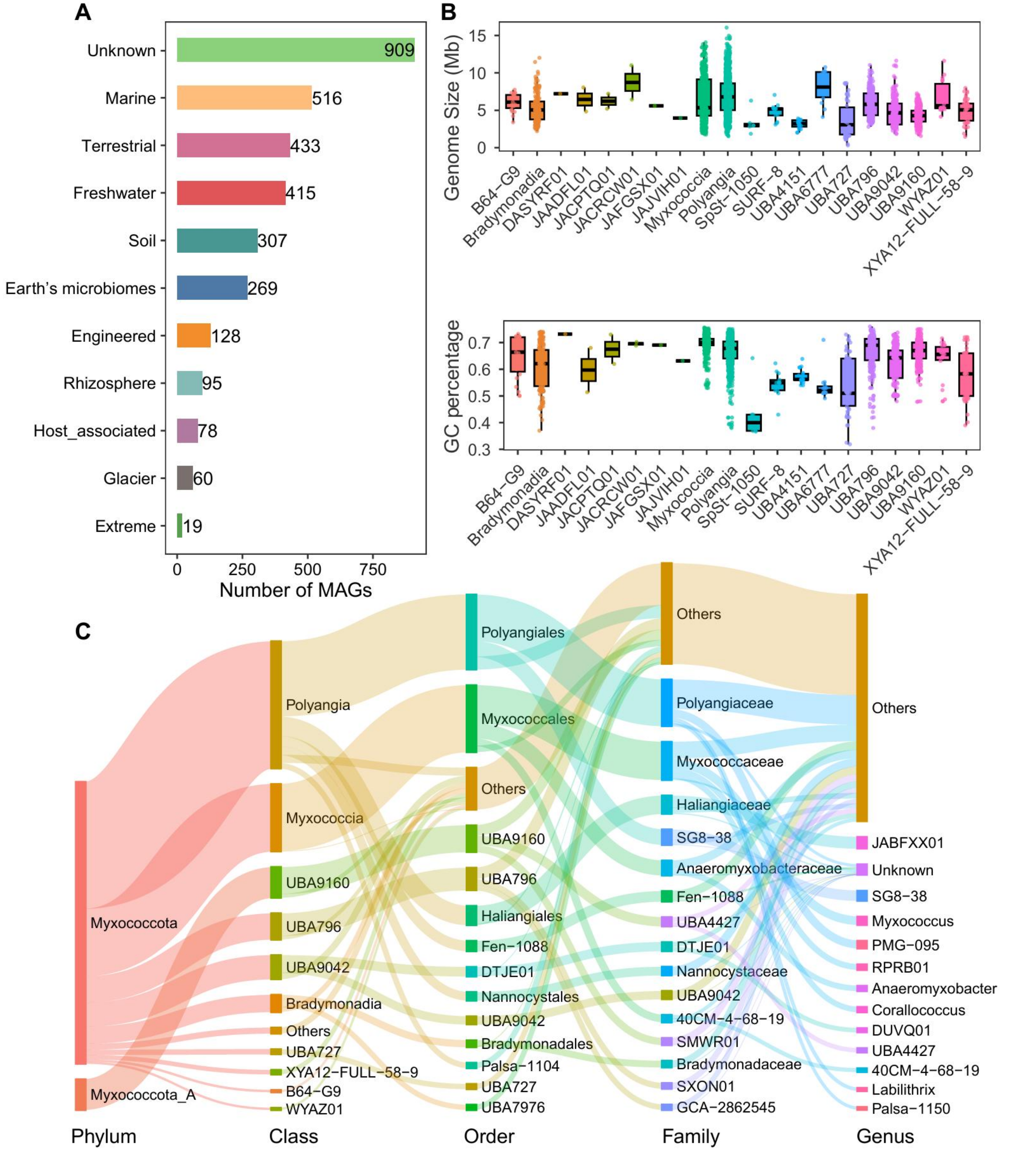


**Fig. S1. Overview of the 3,229 myxobacterial genomes analyzed in this study. (A)** Environmental origins of the 3,229 collected myxobacterial genomes **(B)** Genome size and Guanine-Cytosine (GC) content across myxobacterial classes. **(C)** Taxonomic distribution of the collected genomes at the class, order, and family levels.

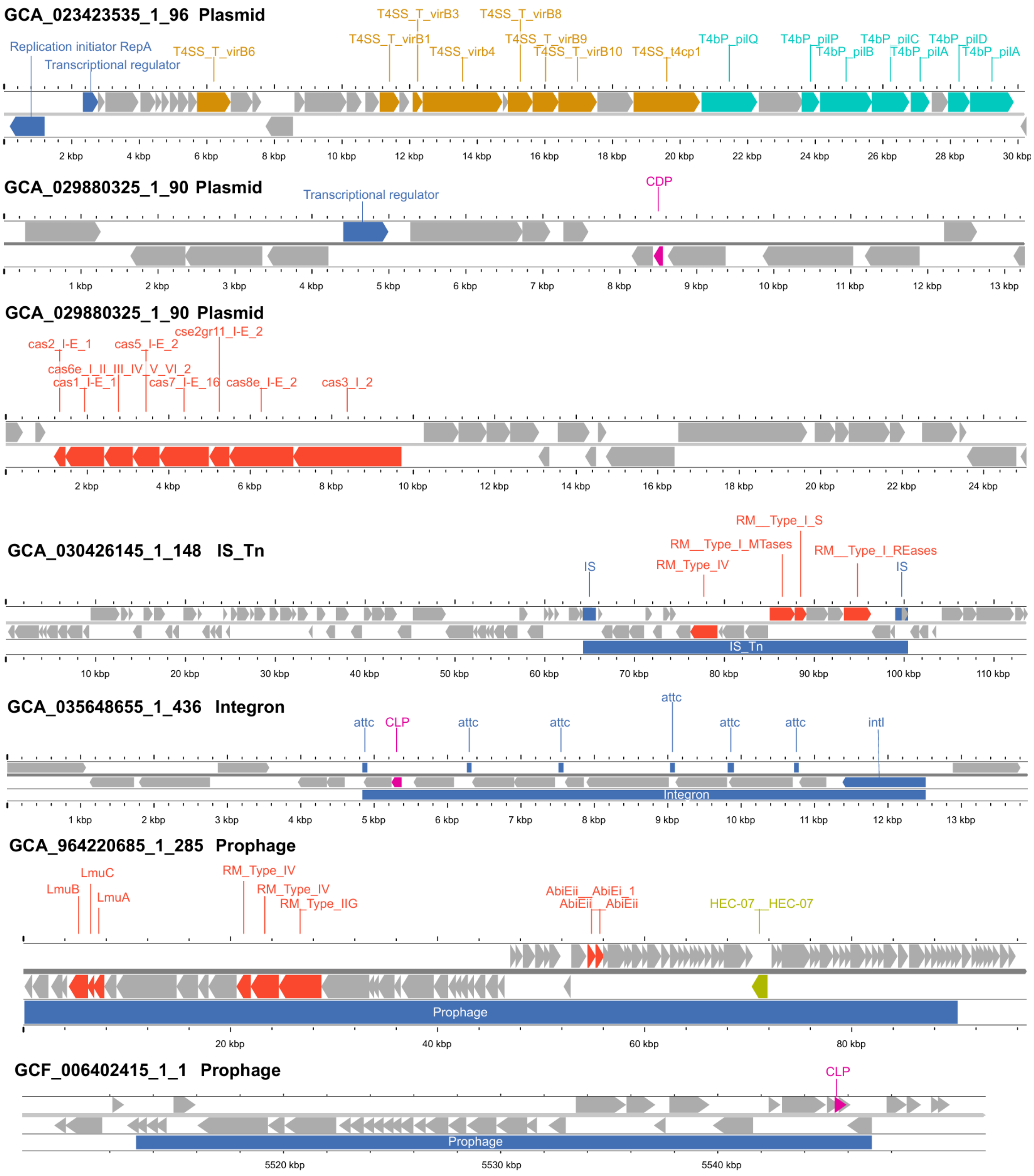


**Fig. S2. Representative mobile genetic elements carrying defense systems, secretion systems, or antimicrobial peptides.** Representative examples of plasmids, composite transposons, integrons, and prophages are shown. Genes are colored according to their predicted functional categories, and the genomic organization of each MGE is displayed to illustrate the genetic context of the associated cargo genes.

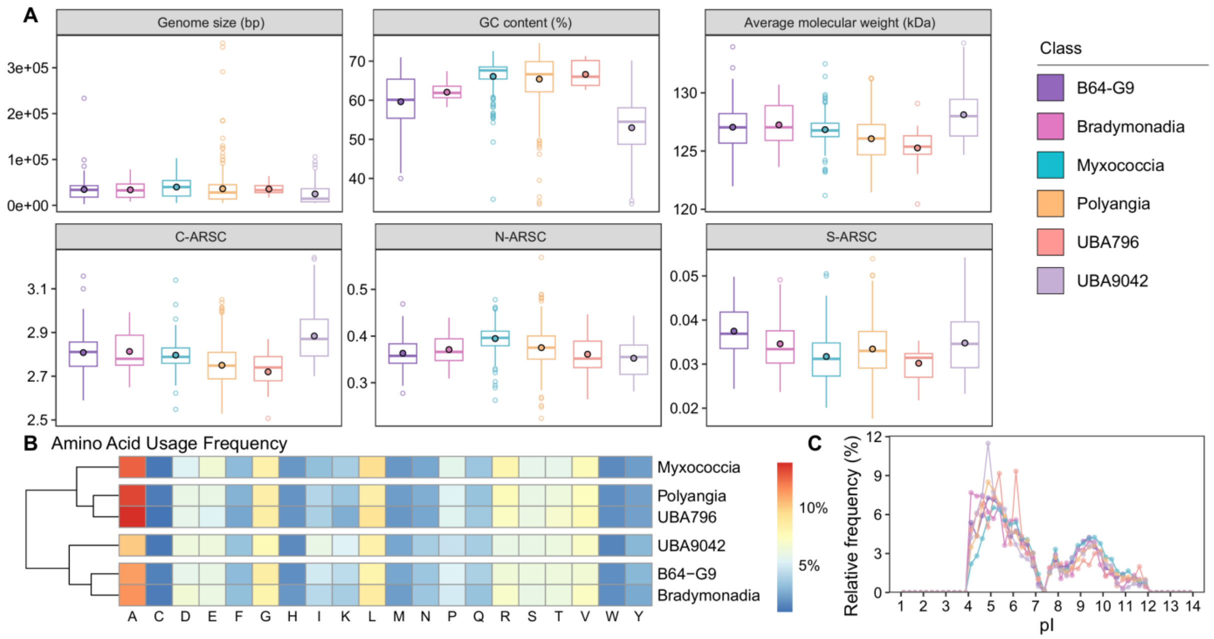


**Fig. S3. Genomic and proteomic characteristics of viruses in the myxobacterial virus database (MVD). (A)** Genome size, GC content, average protein molecular weight, and average number of carbon (C-ARSC), nitrogen (N-ARSC), and sulfur (S-ARSC) atoms per residue side chain (ARSC) of viruses grouped by host class. In each boxplot, the central line and whiskers indicate the median and 1.5 times the interquartile range. The upper and lower sides of boxes represent the interquartile range between 25th and 75th percentile. The dot with black border indicates the mean value. Points beyond whiskers are outliers. **(B)** Heat map showing amino acid usage patterns of viruses grouped by host class. Amino acid frequencies were calculated from all proteins encoded by each viral genome. **(C)** Distribution of predicted protein isoelectric point (pI) values for viruses grouped by host class.

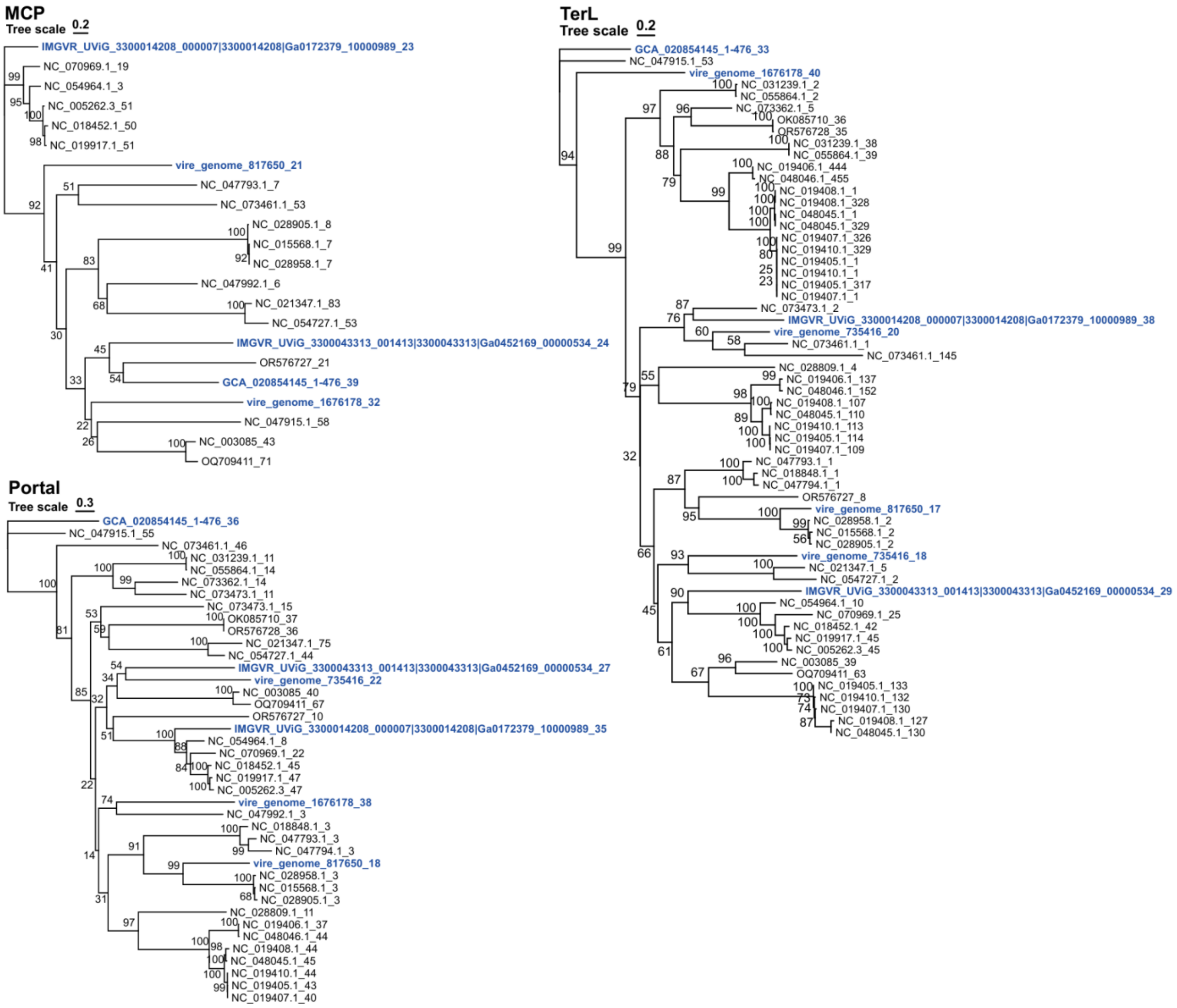


**Fig. S4. Maximum-likelihood phylogenetic tree of major capsid proteins, portal proteins and terminase large subunits from newly proposed viruses and reference viruses.** Bootstrap support values are indicated at internal nodes. Newly proposed viral sequences are highlighted, and reference viral sequences are included for phylogenetic context.

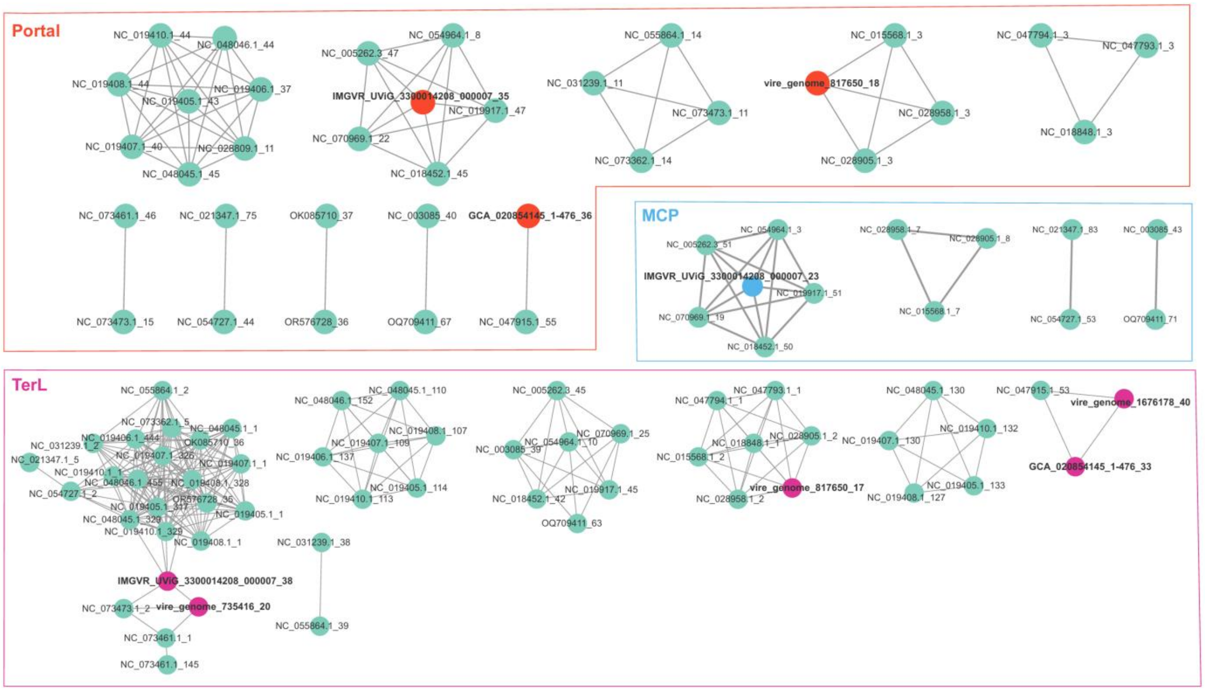


**Fig. S5. Sequence similarity networks reveal distinct clusters of viral structural proteins.** Sequence similarity networks of viral major capsid proteins (MCPs), portal proteins and terminase large subunits (TerLs). Protein sequences are clustered based on pairwise sequence similarity using CLANS. Each node represents a viral protein sequence, and edges indicate significant sequence similarity between protein pairs with a CLANS P value ≤ 0.0001. Distinct protein clusters are indicated by different colors.

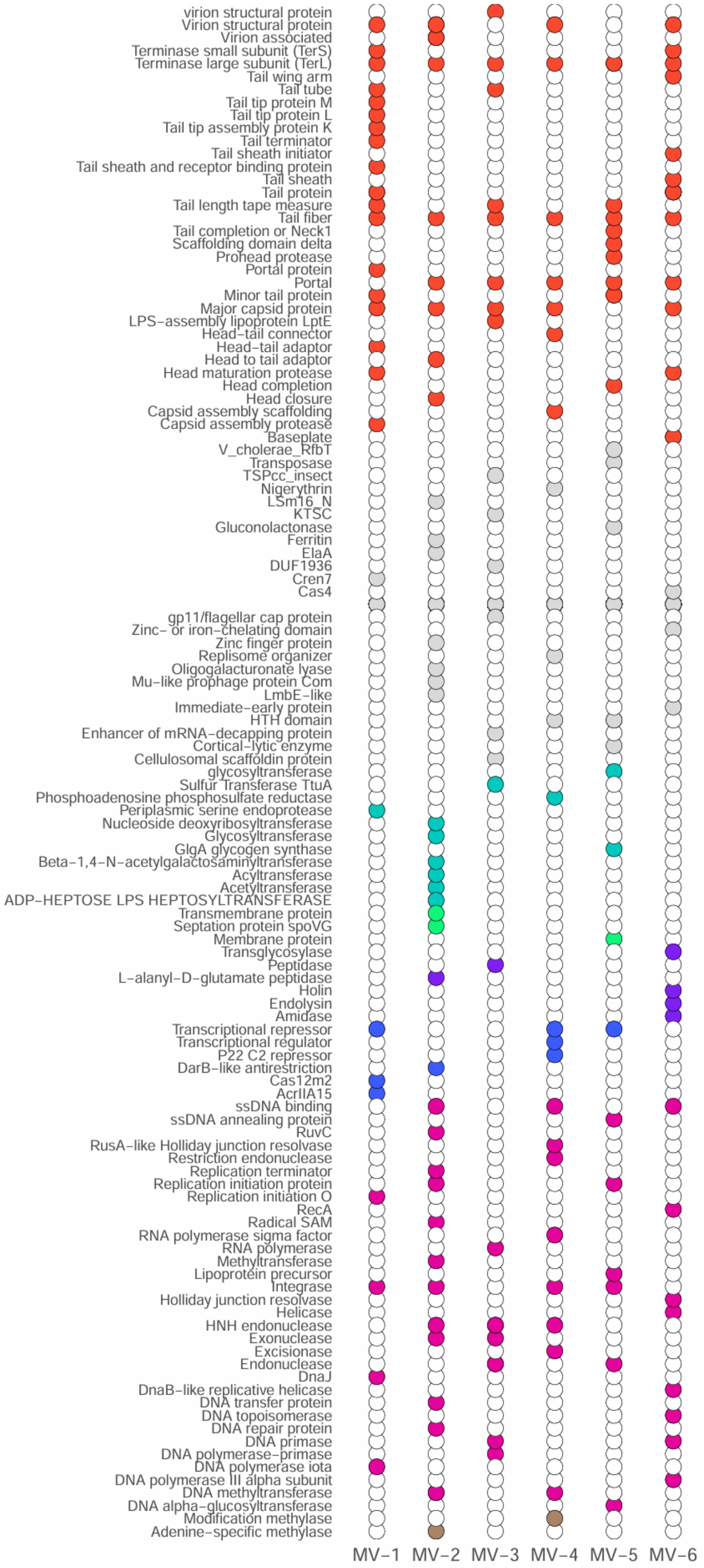
virion structural protein
Virion structural protein
Virion associated
Terminase small subunit (TerS)
Terminase large subunit (TerL)
Tail wing arm
Tail tube
Tail tip protein M
Tail tip protein L
Tail tip assembly protein K
Tail terminator
Tail sheath initiator
Tail sheath and receptor binding protein
Tail sheath
Tail protein
Tail length tape measure
Tail fiber
Tail completion or Neck1
Scaffolding domain delta
Prohead protease
Portal protein
Portal
Minor tail protein
Major capsid protein
LPS−assembly lipoprotein LptE
Head−tail connector
Head−tail adaptor
Head to tail adaptor
Head maturation protease
Head completion
Head closure
Capsid assembly scaffolding
Capsid assembly protease
Baseplate
V_cholerae_RfbT
Transposase
TSPcc_insect
Nigerythrin
LSm16_N
KTSC
Gluconolactonase
Ferritin
ElaA
DUF1936
Cren7
Cas4
gp11/flagellar cap protein
Zinc− or iron−chelating domain
Zinc finger protein
Replisome organizer
Oligogalacturonate lyase
Mu−like prophage protein Com
LmbE−like
Immediate−early protein
HTH domain
Enhancer of mRNA−decapping protein
Cortical−lytic enzyme
Cellulosomal scaffoldin protein
glycosyltransferase
Sulfur Transferase TtuA
Phosphoadenosine phosphosulfate reductase
Periplasmic serine endoprotease
Nucleoside deoxyribosyltransferase
Glycosyltransferase
GlgA glycogen synthase
Beta−1,4−N−acetylgalactosaminyltransferase
Acyltransferase
Acetyltransferase
ADP−HEPTOSE LPS HEPTOSYLTRANSFERASE
Transmembrane protein
Septation protein spoVG
Membrane protein
Transglycosylase
Peptidase
L−alanyl−D−glutamate peptidase
Holin
Endolysin
Amidase
Transcriptional repressor
Transcriptional regulator
P22 C2 repressor
DarB−like antirestriction
Cas12m2
AcrIIA15
ssDNA binding
ssDNA annealing protein
RuvC
RusA−like Holliday junction resolvase
Restriction endonuclease
Replication terminator
Replication initiation protein
Replication initiation O
RecA
Radical SAM
RNA polymerase sigma factor
RNA polymerase
Methyltransferase
Lipoprotein precursor
Integrase
Holliday junction resolvase
Helicase
HNH endonuclease
Exonuclease
Excisionase
Endonuclease
DnaJ
DnaB−like replicative helicase
DNA transfer protein
DNA topoisomerase
DNA repair protein
DNA primase
DNA polymerase−primase
DNA polymerase iota
DNA polymerase III alpha subunit
DNA methyltransferase
DNA alpha−glucosyltransferase
Modification methylase
Adenine−specific methylase
MV−1
MV−2
MV−3
MV−4
MV−5
MV−6

**Fig. S6. Distribution of genes among six viruses.** Distribution of annotated genes among six viruses. Each column represents a virus, and each row represents a gene (lower y axis). Filled circles indicate the presence of an annotated orthologue, whereas open circles indicate the absence of an identified orthologue.

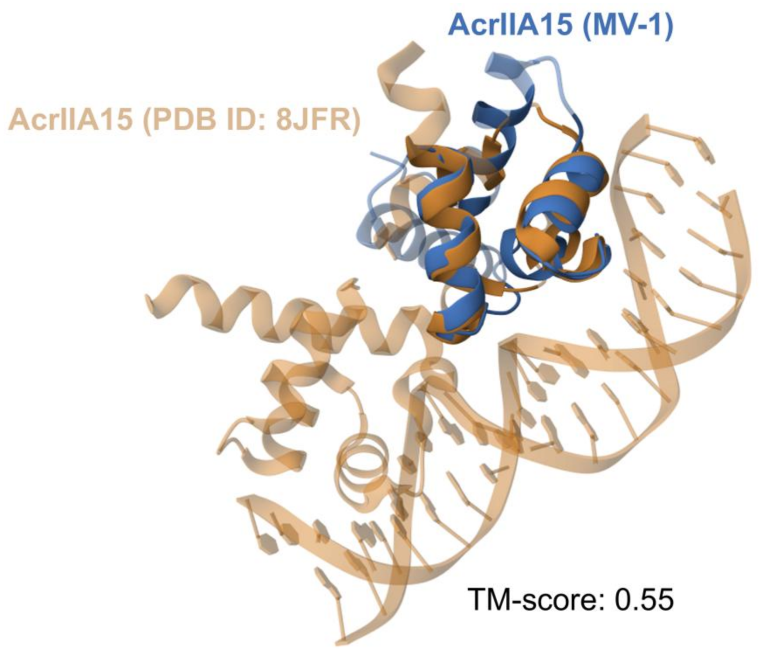


**Fig. S7. Structural similarity between the predicted anti-CRISPR protein and an experimentally validated homolog.** AlphaFold3-predicted structure of the anti-CRISPR (Acr) protein identified in this study. The predicted Acr structure (blue) is superimposed onto an experimentally validated Acr structure (yellow; obtained from the Protein Data Bank (PDB)) using the Matchmaker function in ChimeraX.

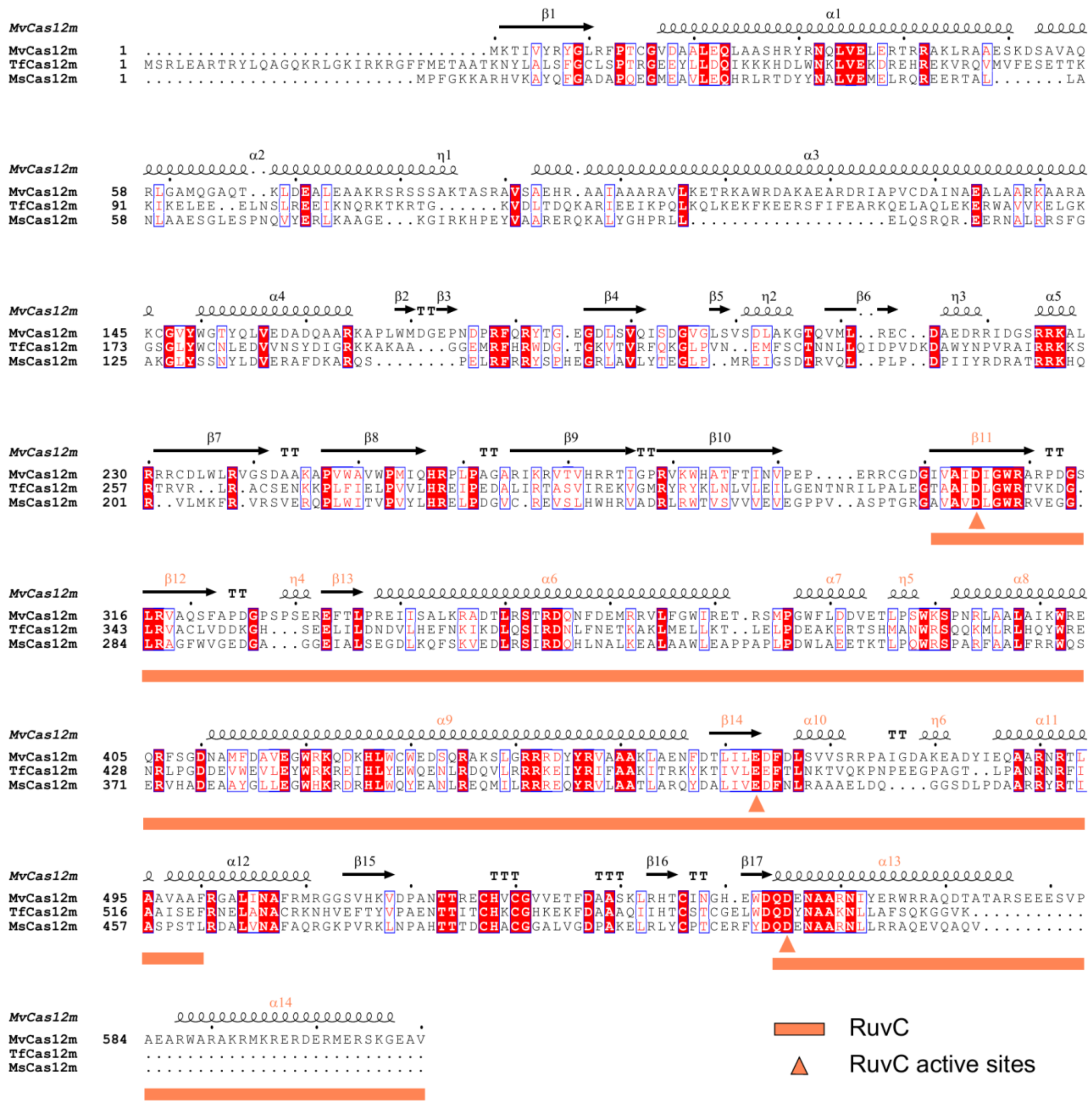


**Fig. S8. Sequence conservation and structural features of Cas12m proteins. Protein alignment of MvCas12m, TfCas12m, MsCas12m.** Multiple sequence alignment of MvCas12m, TfCas12m and MsCas12m proteins. The predicted secondary structure elements of MvCas12m are shown above the alignment. Conserved residues within the RuvC nuclease domain are indicated by triangles below the alignment. The alignment was generated using Clustal Omega and visualized with ESPript3.

### Below tables are provided in spreadsheet format (.xlsx).

**Supplementary Table 1.** Metadata of 3,229 myxobacterial genomes.

**Supplementary Table 2.** Defense systems identified in 3,229 myxobacterial genomes.

**Supplementary Table 3.** Secretion systems identified in 3,229 myxobacterial genomes.

**Supplementary Table 4.** Antimicrobial peptides identified in 3,229 myxobacterial genomes.

**Supplementary Table 5.** Biosynthetic gene clusters identified in 3,229 myxobacterial genomes.

**Supplementary Table 6.** Insertion sequences identified in 3,229 myxobacterial genomes.

**Supplementary Table 7.** Integrons identified in 3,229 myxobacterial genomes.

**Supplementary Table 8.** Plasmids identified in 3,229 myxobacterial genomes.

**Supplementary Table 9.** Phages identified in 3,229 myxobacterial genomes using geNomad.

**Supplementary Table 10.** Integrative and conjugative elements identified in 3,229 myxobacterial genomes.

**Supplementary Table 11.** CRISPR‑Cas systems identified in 3,229 myxobacterial genomes.

**Supplementary Table 12.** Metadata of 790 myxophages.

**Supplementary Table 13.** Metadata of six proposed novel viral groups.

**Supplementary Table 14.** Functional annotation of six newly proposed myxophages.
